# Designed Ankyrin Repeat Proteins provide insights into the structure and function of CagI and are potent inhibitors of CagA translocation by the *Helicobacter pylori* type IV secretion system

**DOI:** 10.1101/2022.11.08.515452

**Authors:** Marine Blanc, Clara Lettl, Jérémy Guérin, Anaïs Vieille, Sven Furler, Sylvie Briand-Schumacher, Birgit Dreier, Célia Bergé, Andreas Plückthun, Sandrine Vadon-Le Goff, Rémi Fronzes, Patricia Rousselle, Wolfgang Fischer, Laurent Terradot

## Abstract

The bacterial human pathogen *Helicobacter pylori* produces a type IV secretion system (*cag*T4SS) to inject the oncoprotein CagA into gastric cells. The *cag*T4SS external pilus mediates attachment of the apparatus to the target cell and the delivery of CagA. While the composition of the pilus is unclear, CagI is present at the surface of the bacterium and required for pilus formation. Here, we have investigated the properties of CagI by an integrative structural biology approach. Using Alpha Fold 2 and Small Angle X-ray scattering, it was found that CagI forms elongated dimers mediated by rod-shaped N-terminal domains (CagI^N^) and prolonged by globular C-terminal domains (CagI^C^). Three Designed Ankyrin Repeat Proteins (DARPins) K2, K5 and K8 selected against CagI interacted with CagI^C^ with subnanomolar affinities. The crystal structures of the CagI:K2 and CagI:K5 complexes were solved and identified the interfaces between the molecules, thereby providing a structural explanation for the difference in affinity between the two binders. Purified CagI and CagI^C^ were found to interact with adenocarcinoma gastric (AGS) cells, induced cell spreading and the interaction was inhibited by K2. The same DARPin inhibited CagA translocation by up to 65% in AGS cells while inhibition levels were 40% and 30% with K8 and K5, respectively. Our study suggests that CagI^C^ plays a key role in *cag*T4SS-mediated CagA translocation and that DARPins targeting CagI represent potent inhibitors of the *cag*T4SS, a crucial risk factor for gastric cancer development.

**Author summary:** *Helicobacter pylori* is a bacterial pathogen that colonises the human stomach in half of the world’s population. The most virulent strains use the cag- type IV secretion system (*cag*T4SS), a molecular nanomachine capable of injecting the oncoprotein CagA into gastric cells. How CagA is delivered is unknown, but the *cag*T4SS produces an external appendage referred to as pilus, which interacts with host cell receptors, mediating CagA translocation from the cytoplasm of the bacteria to the inner membrane of the host cell. In this study we have investigated the structural and functional properties of CagI, a protein long-thought to be associated with the *cag*T4SS pilus but with yet unknown function. We found that CagI displays a unique dimeric structure and that its C-terminal domain is involved in interaction with the host cell. Designed Ankyrin Repeat Proteins were selected against CagI and found to interact with its C-terminal moiety with high affinity. DARPin binding was able to prevent CagI interaction with the host cell and inhibited CagA translocation by *H. pylori*. Our study reveals the role of CagI in *cag*T4SS interaction with gastric cells and provides a first example of a small protein binder inhibiting the *cag*T4SS activity.

## Introduction

*Helicobacter pylori* is a Gram-negative bacterium that colonizes the human stomach in half of the world’s population, and it is a major risk factor for the development of gastric diseases, including ulcers and gastric cancers [1]. Strains carrying the cag-pathogenicity island (*cag*PAI) are more frequently associated with severe diseases [2]. This *cag*PAI is a 40 kbp DNA region that encodes for a type IV secretion system (*cag*T4SS) and for the CagA oncoprotein. Upon contact with gastric cells, the *cag*T4SS delivers CagA into epithelial gastric cells. Once injected, CagA attaches to the inner leaflet of the membrane of the cell where it can be phosphorylated by host cell kinases and interacts with a plethora of cell signalling proteins, hereby promoting tumor development [3]. Although CagA is the only protein effector, several molecules have been reported to be translocated by the *cag*T4SS machinery, including DNA, peptidoglycan, or ADP-heptose that have important pro-inflammatory effects [4].

T4SSs are versatile multi-protein bacterial devices used to transport macromolecules across membranes in various biological processes such as natural transformation, conjugation or delivery of protein effectors into target cells [5, 6]. Based on the prototypical T4SS VirB/D from *Agrobacterium tumefaciens*, T4SSs consist of 12 proteins, VirB1-B11 and VirD4, that form a molecular nanomachine. T4SSs generally comprise a core machinery, made of heteromultimers of VirB3-10 that form a stable complex spanning both bacterial membranes [7]. This core machine is used to produce the external pilus and to translocate substrates [5]. Pilus biogenesis requires the recruitment of VirB11 to VirB4 that promotes the assembly of VirB2 subunits and phospholipids capped by the minor pilin VirB5 [8]. The delivery of substrates relies on the VirD4 ATPase that serves as a coupling protein with the core machinery by interacting with VirB10 [9]. The *cag*PAI encodes for around 27 proteins, some of which are clear homologues of VirB/D proteins but others have no homologues outside *H. pylori* genomes [10]. As a consequence, the core structure of the machinery comprises several additional proteins and is unusually large [4, 11]. Early studies using scanning electron microscopy found that the *cag*T4SS pilus appeared as a flexible sheathed structure protruding outside the cells [12, 13]. Similar tubular structures containing lipopolysaccharides that were located nearby *cag*T4SS complexes were more recently imaged by cryo-electron tomography [14]. Nevertheless, the *cag*T4SS pilus composition is unclear and CagC, the homologue of the major pilin VirB2, was found to be dispensable for pilus formation [15].

The production of a functional pilus requires CagL, CagI and CagH, three proteins specific to the *cag*T4SS that are located on the same operon [16]. CagI, CagL and CagH might interact together, and each of them is required for CagA translocation [16, 17]. Much attention and studies have focused on CagL (reviewed in [18]), since the protein mediates cell attachment *in vitro* and possesses an arginine-glycine-aspartate (RGD) motif involved in integrin binding [19–22]. CagL structures [19, 23-25] revealed a six-helix bundle with a cysteine-clasped loop important for its function [26]. Despite having no clear structural homology, CagL is considered a functional homologue of VirB5 since it shares biophysical properties, it is located at the tip of the pilus and interacts with host cell receptors [19, 27]. However, CagL is not the only Cag protein able to bind to integrins [18] and thus multiple interactions might take place at the host-bacterium interface. CagI interacts with integrin α_5_β_1_ *in vitro* and *in vivo* [28, 29] and is detected at the surface of *H. pylori* cells [16, 30, 31]. CagI might be also directly associated with the *cag*T4SS core-complex assembly, since deletion of the *virB8* homologue *cagV*, *cagX* and *cagY* resulted in CagI instability [31, 32]. The C-terminus of CagI shows sequence conservation with CagL including the two cysteines positioned approximately 100 residues upstream of the C-terminal residues [19] and a C-terminal hexapeptide motif essential for CagA secretion [16]. CagI interacts with CagL *in vitro* [28] and influences its stability *in vivo* [31].

Here, we have used an integrated structural biology approach to gain insights into the structure of CagI. Our data reveal that CagI is a modular protein consisting of an N- and C-terminal domain. The N-terminal domain mediates the protein dimerization and the C-terminal domain is able to mediate cell adhesion and spreading. We also selected Designed Ankyrin Repeat Proteins (DARPins, [33]) against the CagI protein to probe its function. DARPins are small (14 −18 kDa), highly stable, α-helical scaffolds that can bind with high affinity to their targets and have thus applications in various fields, including crystallography, diagnostics and therapeutics [33]. Each repeat consists of 33 amino acids, of which 7 are randomized. Two or three internal repeats are stacked and are flanked by N- and C-terminal capping repeats, to result in N2C or N3C structures. Three DARPins targeting the C-terminal domain of CagI were found to inhibit CagI-mediated cell adhesion and CagA translocation in human cells by up to 65%. These results point towards a key role for CagI in the injection of the main *cag*T4SS effector, possibly by facilitating pilus adhesion to the host cell receptors, thereby identifying a potential way to inhibit cagTSS and help control *H. pylori*-mediated oncogenesis.

## Results

### CagI forms elongated dimers assembled via the N-terminal region

To investigate the structure of CagI, the protein (residues 21 to 381) was purified and its molecular mass determined by size exclusion chromatography coupled to multi-angle light scattering (SEC-MALS). CagI eluted as a single peak with a mass of 79 kDa, consistent with a dimer (Fig. 1A). We then used AlphaFold 2 (AF) [34] to predict the CagI dimer structure and generated three different models (Fig. S1). The monomers of CagI in the three models showed an all α-helical structure consisting of two defined domains (Fig. 1B). The N-terminal region (CagI^N^) comprising residues 21 to 190 consists of two extended helices (α1 and α2) forming a helical hairpin, followed by a short helix α3 (Fig. 1B). α3 connects CagI^N^ to the protein C-terminal domain (CagI^C^) encompassing residues 191 to 381. CagI^C^ is a globular domain made of a four-helix bundle (α4-α7), reminiscent with CagL structure (see below). CagI^C^ models showed some variations in the orientation of α7 but also in the conformation of α6 that was split in two helices with a kink between residues 283 and 286 in one model. CagI structures were predicted with low predicted per-residue confidence score (pLDDT) of 30 to 50, except for residues 205 to 290 that were nearly identical in all models and encompassed helix α3 to half of α6 (Fig. S1).

**Figure 1.**
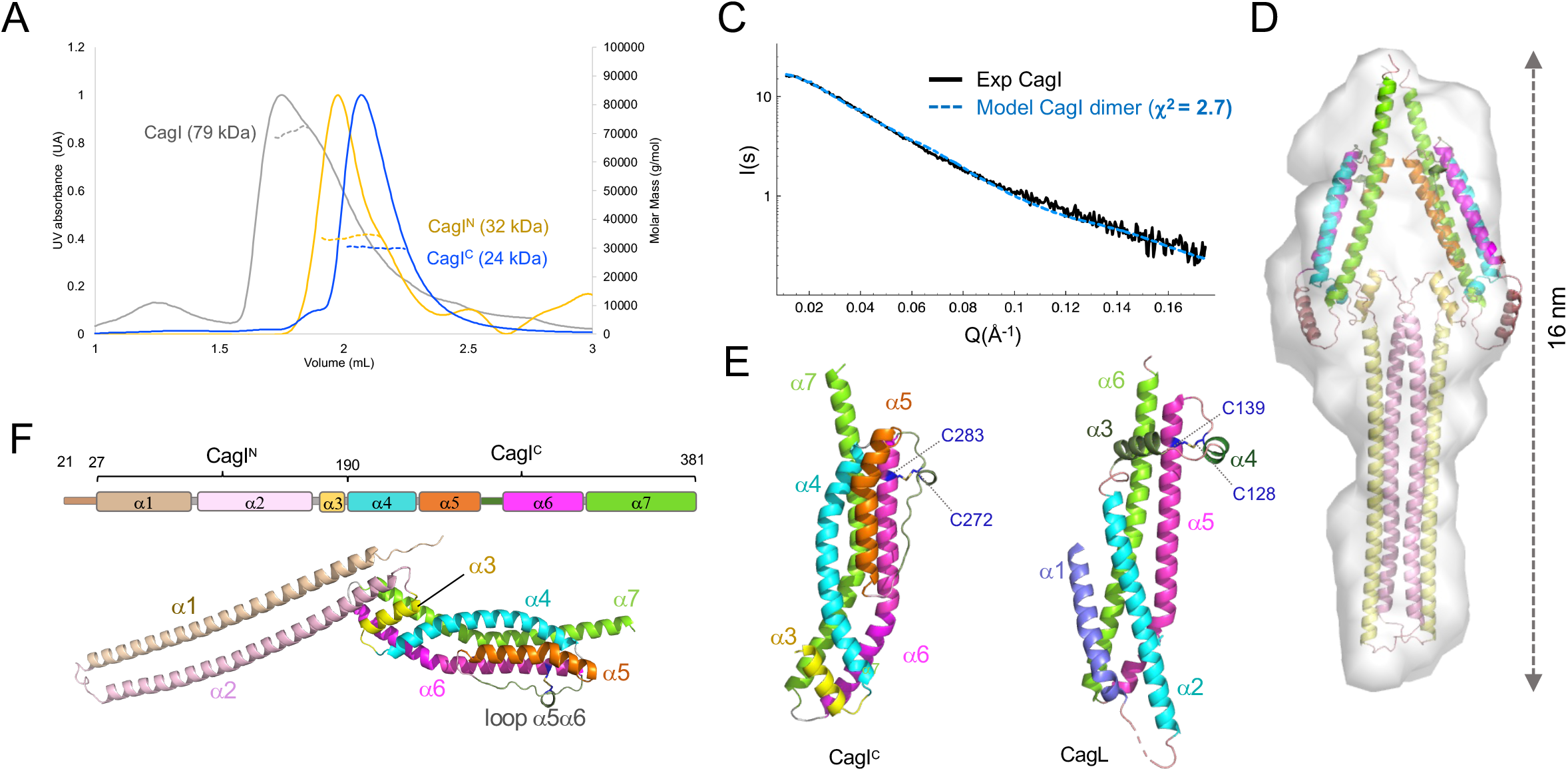
Integrative structural biology study of CagI. A) Size exclusion chromatograms (A_280_) of CagI, CagI^N^ and CagI^C^. MALS weight-averaged molar masses are indicated as dotted lines. B) Schematic representation of CagI predicted secondary structures (top) and cartoon representation of Alpha Fold (AF) model of the CagI monomer with helices coloured as in the schematic view. C) Comparison of CagI dimer theoretical SAXS curve with experimental curve. D) Cartoon depiction of the AF model of CagI dimer coloured as in A) fitted in the SAXS envelope obtained with DAMMIF. E) Comparison of CagI^C^ and CagL (PDB ID: 4YVM) depicted as cartoon. CagI is coloured as in A). CagL secondary structure elements equivalent of those of CagI are coloured accordingly. Cysteine residues involved in disulfide bridges are coloured in dark blue and displayed as ball-and-stick.

AF predicted that the helical hairpin of the N-terminal region was involved in coiled-coil association in two out of three CagI dimer models (Fig. S1). To evaluate the contribution of the N- and C-terminal domains to CagI oligomerisation, we produced them separately and used SEC-MALS to measure their molecular mass (Fig. 1A). CagI^N^ (17 kDa) was found to have a mass of 32 kDa, thus consistent with a dimer in solution. The mass of CagI^C^ (21.5 kDa) measured by MALS was 24 kDa, demonstrating that the isolated domain was monomeric in solution. To gain insight into the overall architecture of the protein in solution, we turned to size exclusion chromatography coupled to small angle X-ray scattering (SEC-SAXS) and collected data on CagI, CagI^N^ and CagI^C^. Comparison of the experimental SAXS curve of CagI with the theoretical ones obtained with AF models showed that model 3 and 2 display χ^2^ values of 2.7 and 3.8, respectively, while model 1 showed the highest value 10.2 (Fig. 1C and Fig. S2). Fitting the dimer model 3 in the *ab initio* DAMMIF envelope confirmed the general shape of the CagI dimer (Fig. 1D). Similar experiments performed with the individual domains CagI^N^ and CagI^C^ confirmed that model 3 fitted better (Fig. S2). Molecular weight calculation using SAXSMoW2 [35] confirmed that CagI^N^ forms a dimer and CagI^C^ is monomeric, in agreement with SEC-MALS data (Table S1). Thus we concluded that, CagI is a dimer in solution mediated by a head-to-head coiled-coil of N-terminal extended α1-α2 hairpins followed by individual CagI^C^ domains.

CagI was proposed to have homology to the other pilus-associated protein CagL based on sequences and motif similarities [25]. Structural superimposition of CagI model with the CagL structure (PDB code 4YVM, [23]) shows that the similarity is limited to CagI helix α6 and the beginning of loop α5-α6 with helix α5 and loop α4-α5 in CagL, respectively (Fig. 1E). This includes a conserved disulfide bridge between C272 and C283 predicted in CagI that tethers the α5-α6 loop to α6, corresponding to C128 and C139 in CagL (Fig. 1E, Fig. S3). The remaining parts of CagL structure and CagI^C^ are rather different with, notably the two short helices α3 and α4 of CagL absent in the CagI structure (Fig. 1B, E, Fig. S3) and an additional helix α5 in CagI.

### DARPins against *Helicobacter pylori* CagI bind with high affinity to the C-terminal domain of the protein

To generate DARPin binders against CagI, avi-tagged CagI protein (CagI_avi_) was immobilized alternatingly on streptavidin and neutravidin. Ribosome display selections were performed over four round using a semi-automated 96-well plate format (see Materials and Methods). After the fourth round of ribosome display selection, 380 single DARPin clones were expressed with N-terminal MRGS(H)_8_ and C-terminal FLAG tag and screened for binding to CagI with a C-terminal strep-tag (CagI_strep_) using ELISA on streptactin-coated plates. From the initial hits of the 380 analyzed DARPin clones, 32 were randomly chosen for further analysis and their sequence determined. Of these, 15 were identified as unique clones and they were expressed in a 96-well format and IMAC purified. The purified DARPins were used in a hit validation to bind immobilized CagI_strep_ by ELISA using FLAG detection.

Next, we investigated if co-expression of the DARPins with CagI in *E. coli* cells could lead to co-purification. The vector expressing CagI_strep_ was co-transformed in *E. coli* cells with each vector expressing a DARPin (K1 to K15) fused to an N-terminal His_8_-tag. To monitor complex formation, we purified cell extracts on Ni-NTA beads and determined by SDS-PAGE if CagI_strep_ co-purified with the DARPin. We observed that CagI_strep_ co-eluted with DARPin K2, K5, K8, K9, K10, K11, K12 and K15 (Fig. S4). The presence of CagI_strep_ was assessed by Western blotling using antibodies against the strep-tag (Fig. S4) in all elution fractions that were positive in SDS-PAGE. A band corresponding to CagI_strep_ was also detected in the elution fraction of CagI_strep_/K1 although no band was visible in the SDS-PAGE. The amino-acid sequences of the confirmed DARPins are shown in Fig. S5.

We then purified His8-tagged DARPins K2, K5, K8-K12 and K15 and confirmed that they could also bind CagI_strep_ in pull down assays on Ni-NTA beads (Fig. 2A). The same assays performed with individual CagI domains showed that all DARPins interacted with CagI^C^ but not with the CagI^N^ (Fig. 2A). To determine the affinity of the binders for their target, surface plasmon resonance (SPR) experiments were performed by immobilizing CagI or CagI^C^ on the chip and injecting increasing concentrations of DARPins. DARPins showed different binding modes and K_D_ ranges for CagI and CagI^C^ (Fig. 2B, C, Fig. S6 and Table 1). For measurements on full-length CagI, best fits were obtained with the heterogenous ligand interaction model (Fig. S6A), while interactions of DARPins with CagI^C^ fit well the 1:1 binding model (Fig. 2C, Fig. 6B). This suggests that DARPins interact with the C-terminal domain of CagI in a 1:1 manner, and that CagI dimerization reduces the accessibility of the two C-terminal domains. Because K_D_ values for CagI-DARPins measurements were similar between 1:1 binding and heterogenous ligand models, we considered K_D_ values obtained using the 1:1 binding model to compare the affinities between CagI and CagI^C^ (Table 1). The measured K_D_’s indicated that DARPins had high affinity for CagI in the range of 1-10 nM except for K15 whose K_D_ was 73.7 nM. DARPins K2 and K8 had particularly low k_off_, resulting in a strong affinity for CagI. For all the DARPins, affinities for CagI^C^ were ten times higher, in the range of 0.2-1 nM, while for K2, K8 and K11 K_D_’s as low as 0.03 nM, 0.06 nM and 0.09 nM, respectively, were obtained. K15 again showed a weaker K_D_ of 4.89 nM (Fig. S6, Table 1). Interestingly, sequences of DARPin K2 and K5 differed only by a single amino-acid at position 125, being a leucine for K5 and a phenylalanine for K2 suggesting that the binding modes of the two DARPins were similar but with a 10-fold increase in affinity for K2 compared to K5.

**Figure 2.**
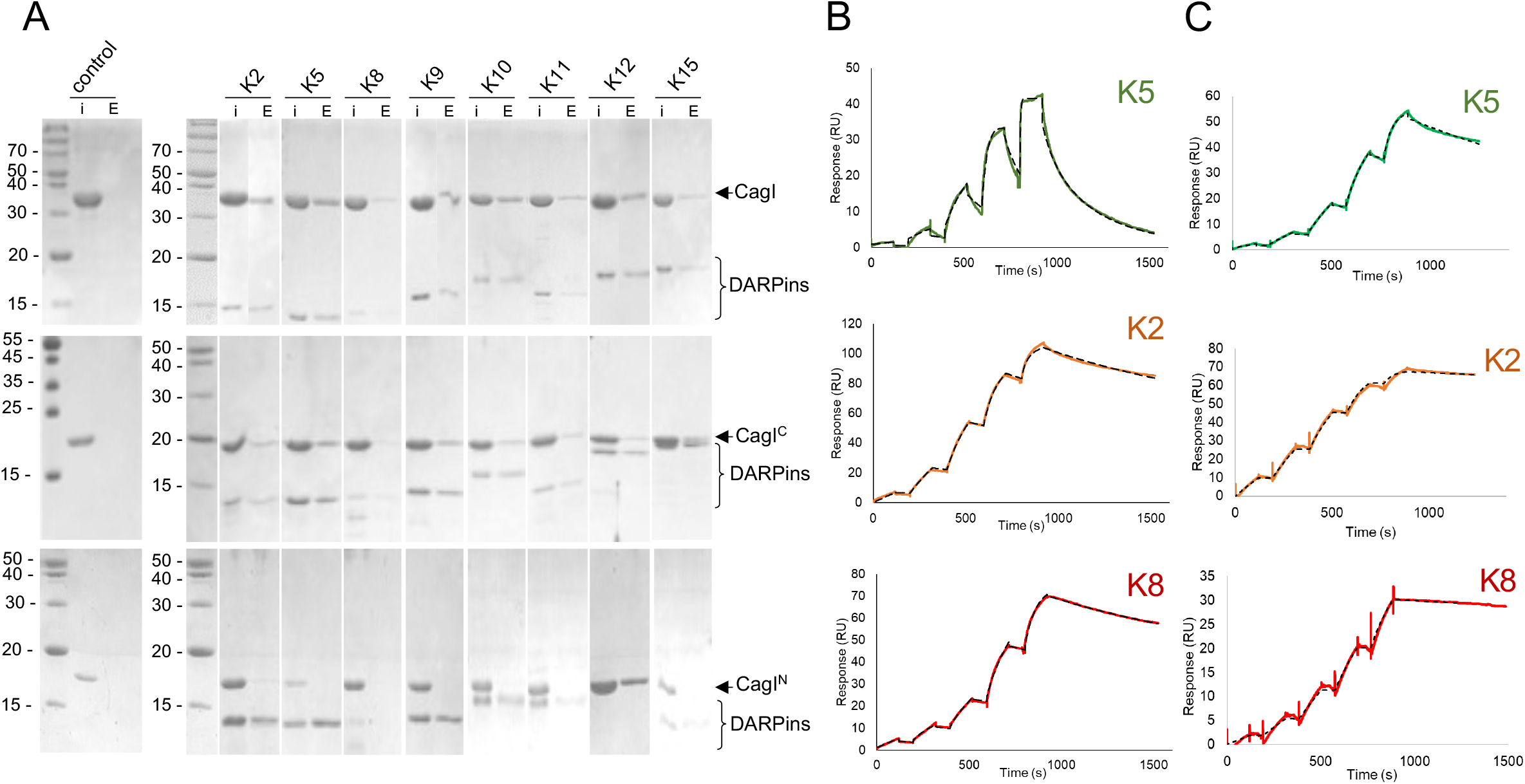
DARPin interaction with CagI. A) Pull down assays of purified untagged CagI (top panels), CagI^C^ (middle panels) or CagI^N^ (bottom panels) with NTA bead-immobilized His_8_-tagged DARPins. “I” denotes input protein and “E” denotes elution. In control experiments proteins were mixed with the resin in the absence of DARPin and were not detected in the elution fraction. B) Representative SPR experiments using single-cycle injection mode on CM5 chips coated with CagI or with C) CagI^C^. DARPins K5 (green curves), K2 (orange curves) or K8 (red curves) were injected on the chips at increasing concentrations as follows. For full-length CagI experiments, concentrations were 0.5, 2.5, 12.5, 62.5 and 312.5 nM for K5 and K8. For K2, concentrations were 1, 3, 9, 27 and 81 nM. For CagI^C^ experiments, K2 and K8 were injected at 0.05, 0.15, 0.45, 1.35 and 4 nM. For K5 concentrations used were 0.11, 0.33, 1, 3 and 9 nM. Fit curves obtained with binding model 1:1 are shown as dashed lines.

**Table 1.** Dissociation constant (K_D_) expressed in nM obtained in Surface Plasmon Resonance experiments with immobilized CagI or CagI^C^ and the DARPins as analytes. Values were obtained using the 1:1 binding model (mean of two separate experiments except for values labeled with a *, for which a single multi-injection experiment was performed).

|  | **CagI** |  |  |  | **CagI<sup>C</sup>** |  |  |  |
| --- | --- | --- | --- | --- | --- | --- | --- | --- |
|  | **$K_D$ (nM)** | **$k_{on}$ (M<sup>-1</sup>s<sup>-1</sup>)** | **$k_{off}$ (s<sup>-1</sup>)** | **$\chi^2$** | **$K_D$ (nM)** | **$k_{on}$ (M<sup>-1</sup>s<sup>-1</sup>)** | **$k_{off}$ (s<sup>-1</sup>)** | **$\chi^2$** |
| K2 | 1.59 ± 0.91 | 1·10<sup>5</sup> | 4·10<sup>-4</sup> | 3-9 | 0.03 ± 0.01 | 2·10<sup>6</sup> | 2·10<sup>-4</sup> | 0.05-1 |
| K8 | 1.15 ± 0.21 | 1·10<sup>5</sup> | 2·10<sup>-4</sup> | 1-10 | 0.06 ± 0.001 | 1·10<sup>6</sup> | 8·10<sup>-5</sup> | 0.05-0.3 |
| K11 | 1.15 ± 0.02 | 8·10<sup>4</sup> | 1·10<sup>-3</sup> | 25-75 | 0.09 ± 0.01 | 4·10<sup>6</sup> | 4·10<sup>-4</sup> | 3.4-4.6 |
| K10 | 3.83 ± 0.37 | 8·10<sup>4</sup> | 5·10<sup>-3</sup> | 9-62 | 0.41 ± 0.10 | 5·10<sup>6</sup> | 6·10<sup>-4</sup> | 2.5-6.3 |
| K9 | 2.80* | 2·10<sup>6</sup> | 4·10<sup>-3</sup> | 44 | 0.22 ± 0.01 | 2·10<sup>6</sup> | 6·10<sup>-4</sup> | 0.3-1.2 |
| K5 | 8.05 * | 2·10<sup>5</sup> | 1·10<sup>-3</sup> | 0.8 | 0.23 ± 0.01 | 2·10<sup>6</sup> | 5·10<sup>-4</sup> | 0.2-1.7 |
| K12 | 8.61 * | 2·10<sup>5</sup> | 1·10<sup>-3</sup> | 43 | 0.81 ± 0.34 | 1·10<sup>6</sup> | 9·10<sup>-4</sup> | 1.2-1.7 |
| K15 | 73.7* | 3·10<sup>4</sup> | 2·10<sup>-3</sup> | 30 | 4.89 ± 2.99 | 5·10<sup>5</sup> | 1·10<sup>-3</sup> | 0.8-4 |

### Structures of CagI:K5 and CagI:K2 complexes

To better understand the molecular basis of DARPin interaction with CagI, we solved the crystal structures of the CagI:K2 and CagI:K5 complexes. The two crystals had very similar cell parameters (see Table 2 for data collection and refinement statistics). The two complex structures revealed that each asymmetric unit contained one DARPin molecule and a fragment of CagI that had undergone proteolysis during crystallisation (Fig. 3A). Density was clear for residues 204-307 of CagI in the two structures and these were nearly identical, with a rmsd of 0.2 Å^2^. The structure of the fragment of CagI consists of a three-helix bundle corresponding to α4, α5 and α6 and the extended α5-α6 loop containing a 3_10_ helix in the AF model (Fig. 3A). Interestingly, the fragment of the CagI crystallised corresponds to the region of the AF model that showed the highest prediction scores. Comparison of the structure of CagI^205-304^ (from the K5:CagI complex) with AF model monomer showed that the prediction was indeed remarkably correct, with a rmsd of 0.9 Å for 103 Cα(Fig. S7). The K5 and K2 interaction site of CagI consists of a hydrophobic groove formed by α4 and α5 residues. Interactions between DARPins and CagI are widespread along the groove and extend to the concave face formed by the DARPin variable loops that wrap around α5 (Fig. 3B). While the interface relies mostly on hydrophobic interactions, five hydrogen bonds also exist between CagI and DARPin residues, respectively: T235 - D112, S242 - D79, E210 - Q28, S242 - H50 and S254 - E22 (Fig. 3B). A single residue difference between K2 and K5 in the C-cap moiety of the DARPin generates small but significant changes in the DARPin/CagI interface. At position 125 the, K5 residue is a leucine and its side chain is placed near a pocket formed by CagI α4 residues T235 and L239 and α5 residues L228 and K225 (Fig. 3C). In K2, the phenylalanine 125 inserts deeper into the pocket and is stabilised by a T-shaped pi-stacking with the nearby Y91. In addition, the side chain of CagI residue K225 makes two hydrogen bonds with the carbonyl of the F125 main chain and the N127 side chain of DARPin K2 (Fig. 3C). As a consequence, the surface buried by complex formation is slightly increased in CagI:K2 with 994 Å^2^ compared to 974 Å^2^ in CagI:K5.

**Table 2.**
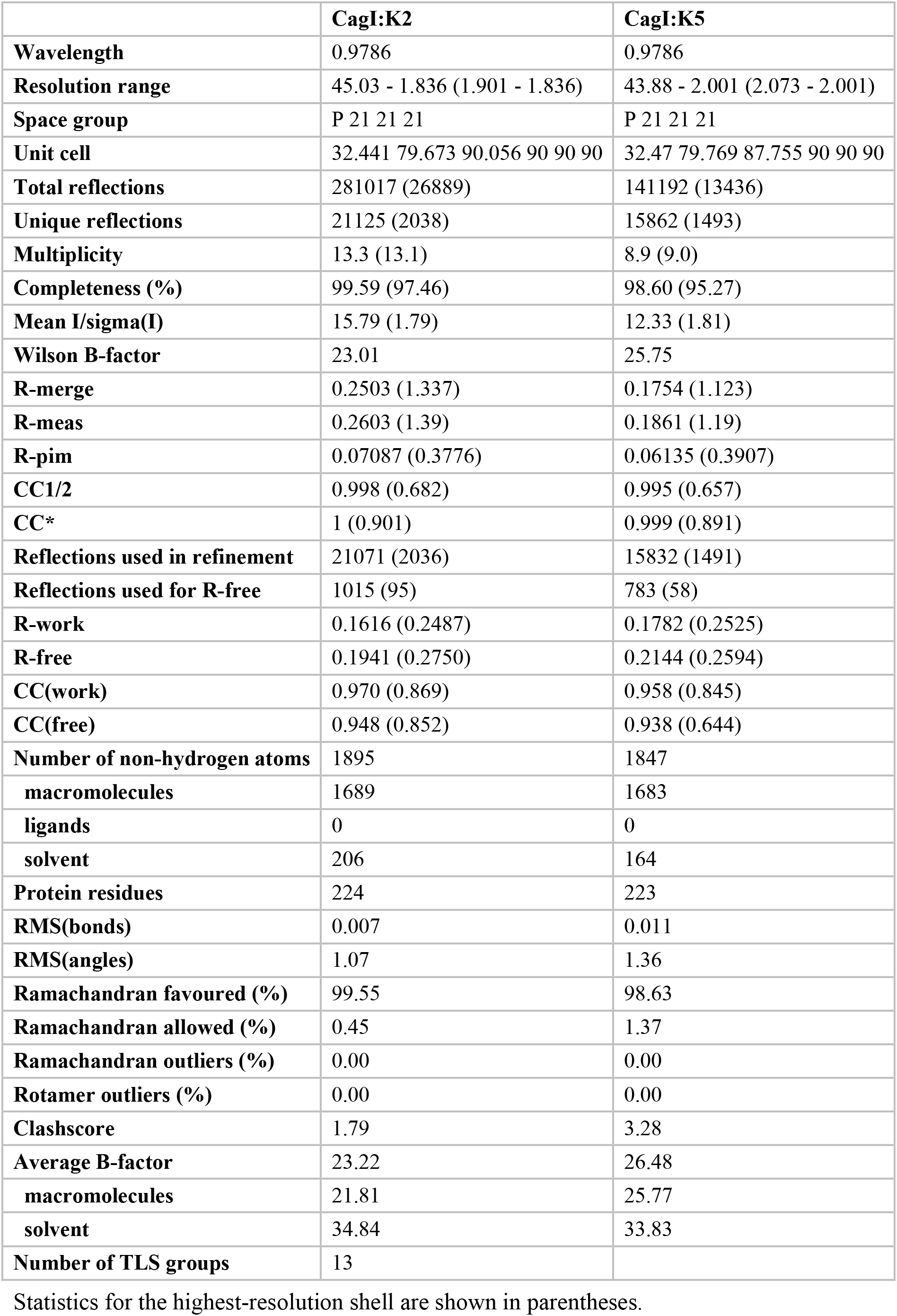
Data collection and refinement statistics

**Figure 3.**
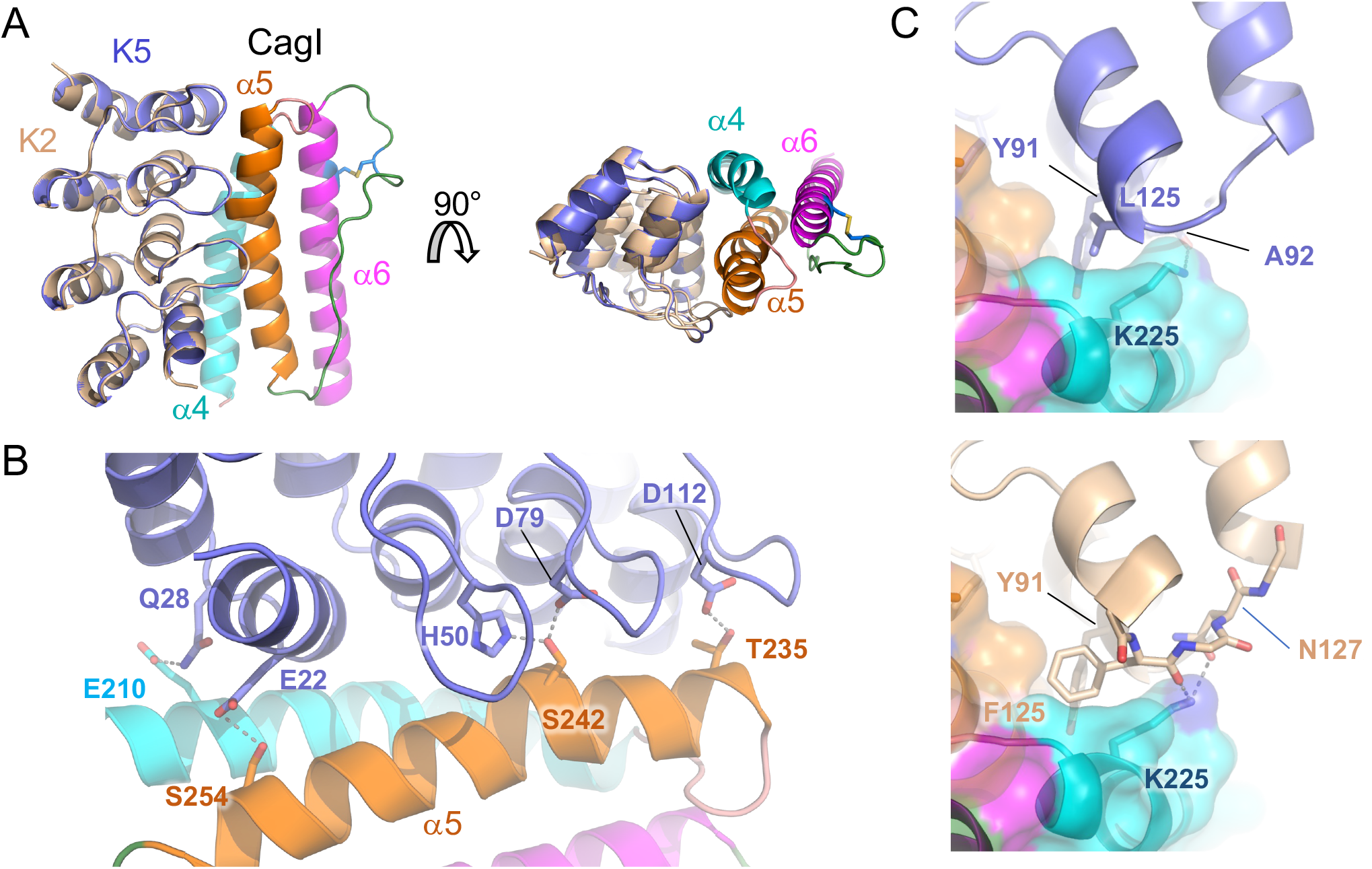
Structures of CagI/DARPin complexes. A) Overview of the structure of CagI:K2 and CagI:K5 complexes. The two structures of DARPin K2 (wheat) and K5 (slate) have been superimposed and are depicted as cartoons. The CagI molecule is displayed as cartoon and surface coloured according to secondary structure (α4 in cyan, α5 in orange and α6 in magenta). Side chains of cysteines 272 and 283 involved in disulfide bridges are shown as ball-and-stick with atoms coloured blue (carbons) and yellow (sulfur). For clarity, only the CagI molecule from the CagI:K2 complex is displayed. B) Detailed view of the interface between DARPin K5 loops and CagI α5 with side chains involved in hydrogen bonds shown as ball-and-stick with atoms coloured as follows: nitrogen in blue, oxygen in red, carbon coloured as in A). Grey dashed lines indicate hydrogen bonds. C) Structural comparison of the CagI:K5 (top) and CagI:K2 (bottom) interface at the groove formed between α4 and α5. Grey dashed lines indicate hydrogen bonds. Detailed view of the interface involving interactions between K2 and CagI helix α4. Close-up view of the interface of CagI:K2 (top panel) and CagI:K5 (bottom panel) showing residues F125 in K2 and L125 in K5 binding to the CagI groove.

### DARPins K2 and K5 interact with two CagI molecules

Analysis of the crystal packing indicates a second interaction site in the CagI:K2 and CagI:K5 structures with significant scores in PISA [36], burying around 800 Å^2^. In addition to the asymmetric unit CagI, K2 and K5 interact with a symmetry-related CagI fragment (noted CagI’). The interface named region 2 (region 1 being the first interface described above) covers CagI’ α4 and part of α6 and involves the convex face of the three variables loops of the DARPins (Fig. 4A). The interactions in that region are the same in CagI:K2 and CagI:K5 complexes. The interface relies on hydrophobic interactions at the groove formed by α4 and α6 and several hydrogen bonds and salt bridges between DARPins D47 and CagI K206 and K213 (Fig. 4B). We therefore evaluated by SEC-MALS the stoichiometry of all CagI/DARPin complexes. CagI:K2, CagI:K5, CagI:K10, and CagI:K8 had a mass of 93-95 kDa, and CagI:K11 had a mass of 90 kDa, all consistent with a monomer of DARPin in complex with a dimer of CagI (Fig. 4C and Fig. S8). CagI:K9 and CagI:K12 calculated masses were 117 kDa and 123 kDa, respectively, consistent with 2 DARPins/2 CagI complexes. No mass could be calculated with CagI:K15 given that the complex was aggregated. Thus we concluded that K2, K5, K8, K10 and K11 were able to bind simultaneously to two CagI^C^ molecules while K9 and K12 bind a single CagI^C^.

**Figure 4.**
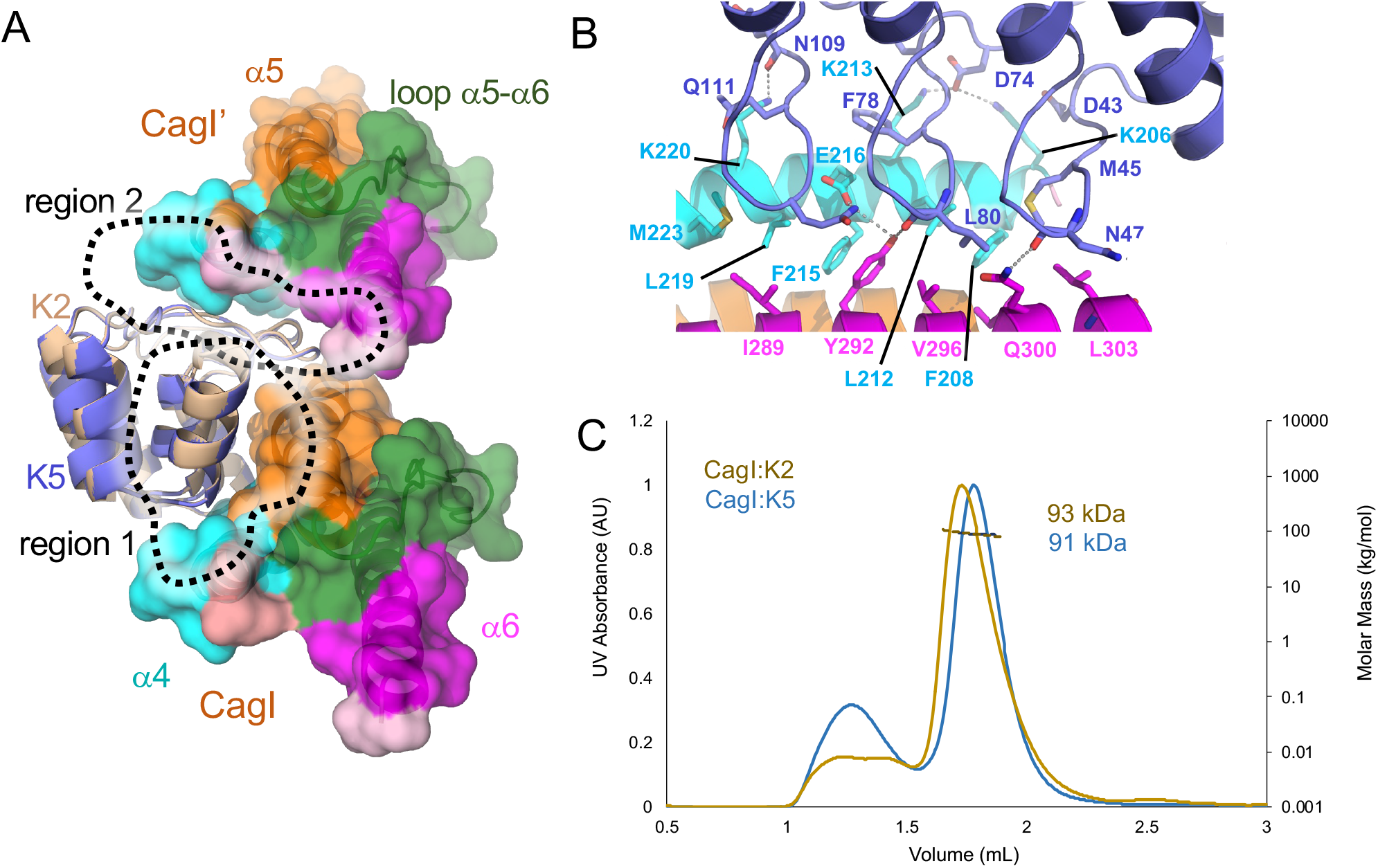
Structural basis for higher affinity of DARPin K2 on CagI. A) Overview of the in the structures of the CagI:K2 and CagI:K5 complexes interfaces. The two DARPin structures K2 (wheat) and K5 (blue) have been superimposed and are depicted as cartoons. The CagI molecule and symmetry related CagI’ are displayed as cartoons and surfaces are coloured as in Fig. 3. For clarity, only CagI molecules from the CagI:K2 complex are displayed. B) detailed view of region 2 interactions between DARPin K5 loop residues and CagI α4 and α6 with involved side chains displayed as ball and sticks with atoms coloured as follows: nitrogen in blue, oxygen in red, carbon as in A). Dashed lines indicate hydrogen bonds. C) SEC-MALS measurements of the CagI:K2 and CagI:K5 purified complexes.

### DARPins targeting CagI inhibit CagA translocation in human adenocarcinoma gastric cells

To determine if the DARPins have an effect on CagA translocation by the *cag*T4SS, we next used a β-lactamase (TEM-1) assay as a reporter system for CagA type IV secretion into gastric epithelial cells [37]. Bacteria producing a TEM-1-CagA fusion were co-incubated with a gastric adenocarcinoma cell line (AGS) in the presence or absence of DARPins. Translocated β-lactamase hydrolyses a fluorophore in the AGS cells and translocation of TEM-1-CagA was thus determined via blue-to-green fluorescence ratios. We observed that bacteria incubated with DARPins K9, K10, K11, K12 and K15 translocated CagA as efficiently as in the control experiment (Fig. 5A). In contrast, a 30% reduction was seen with bacteria incubated with K5, and even a reduction by 40% and 60% in CagA translocation after preincubation with K8 and K2, respectively (Fig. 5A). Similar results were obtained with a HiBiT-CagA translocation assay, which uses a split-luciferase system as a translocation reporter [38]. In this assay, pretreatment of P12[HiBiT-CagA] with DARPins K5, K8 and K2 also resulted in a decrease of translocation levels to comparable levels of the TEM-1-CagA translocation assay (even though K8 did not reach statistical significance), whereas treatment with K9 did not decrease translocation efficiency (Fig. 5B). To determine if these effects were due to a lack of attachment of the whole bacteria to the AGS cells, we monitored bacterial adhesion via flow cytometry using *H. pylori* cells expressing GFP. The results show no major differences in cell adhesion (Fig. S9), suggesting that the DARPins did not affect *H. pylori* binding to AGS cells.

**Figure 5.**
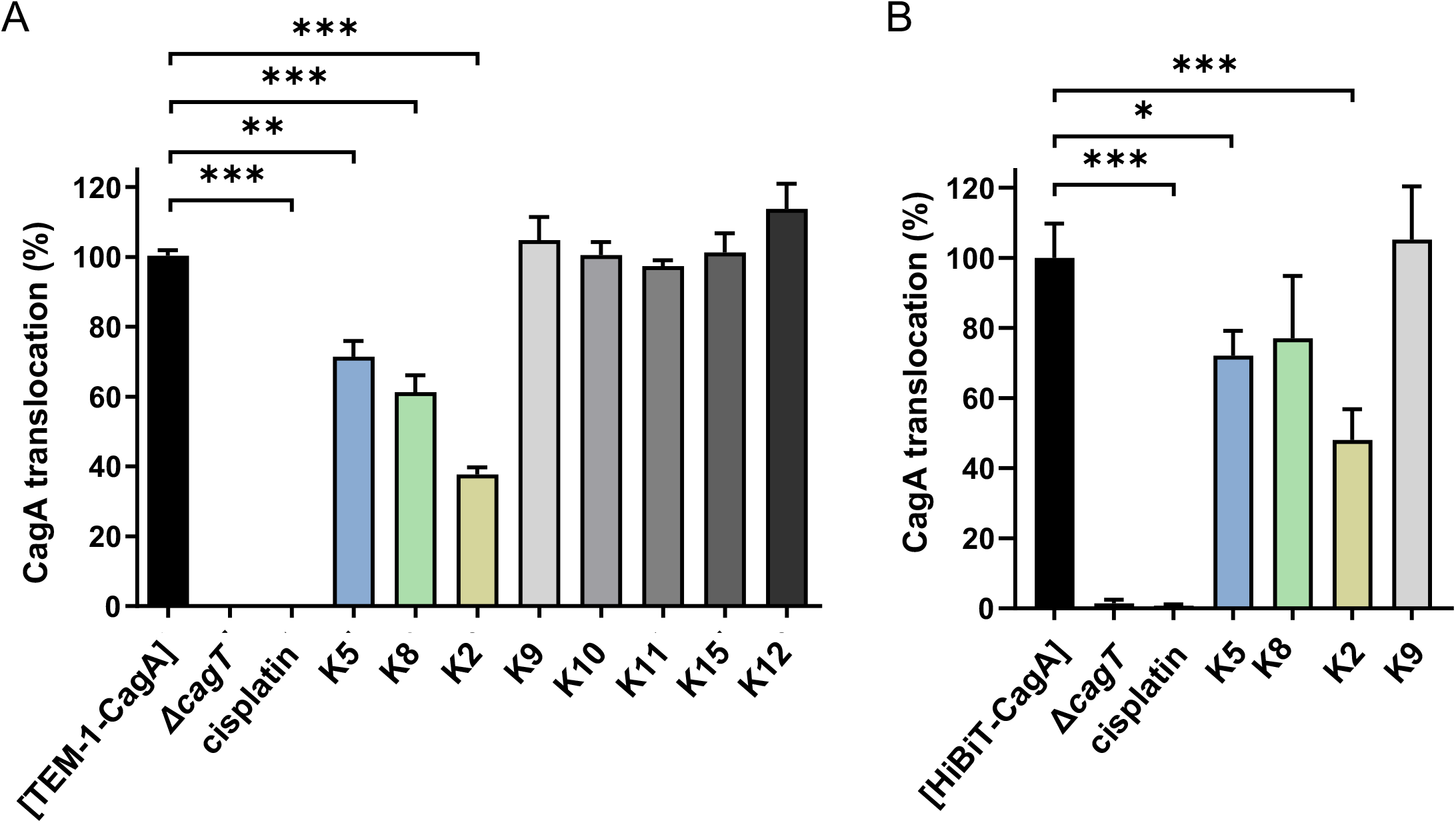
CagA translocation inhibition by DARPins. A) *H. pylori* P12 [TEM-1-CagA] was co-incubated for 2.5 h with AGS cells in the absence or presence of the indicated DARPins at a concentration of 5 μM, and CagA translocation was determined by a TEM-1-CagA translocation assay. As controls, the secretion-deficient mutant P12Δ*cagT* [TEM-1-CagA] was used without pre-treatment, or P12 [TEM-1-CagA] was pre-incubated for 30 min with 100 μM cisplatin. Data are indicated in relation to untreated control, which was set to 100%, and they represent mean values and standard deviations of five independent experiments. (One-way ANOVA; Tukey post-hoc test; **, p<0.01; ***, p<0.001). B) *H. pylori* P12 [HiBiT-CagA] was either left untreated or pre-treated for 30 min with 5 μM of the indicated DARPins or 100 μM cisplatin in PBS/10 % FCS at 37 °C, 10% CO_2_, and the bacterial suspension were used to infect AGS [LgBiT] cells for 2.5 h. Luminescence values were recorded and normalized to untreated control. Data are indicated as mean values with standard deviations resulting from four independent experiments. (One-way ANOVA; Tukey post-hoc test; *, p<0.05; ***, p<0.001).

### CagI mediates cell binding and spreading via its C-terminal domain

Previous studies showed that CagL mediates AGS cell adhesion *in vitro* in a manner reminiscent of fibronectin [20]. Given that CagI and CagL share structural homology, we investigated if CagI could also have a similar effect on human cells. An assay was implemented to compare the binding of AGS cells to purified CagL, CagI or CagI N and C-terminal domains. Multiwell plates were first coated with different amounts of the proteins and then incubated with AGS cells. The extent of AGS cell adhesion was measured 60 min after cell seeding using a colorimetric reaction as described in Materials and Methods. While CagI^N^ did not induce any cellular adhesion, AGS cells adhered to the three other substrates in variable proportions. CagL and CagI^C^ induced the strongest cellular adhesion (Fig. 6A). A variation of the cellular morphology could be noticed according to the ligands (Fig. 6B).While the majority of the cells remain rounded or slightly spread out after both CagI and CagL binding, all the cells appear fully spread on CagI^C^, suggesting a rapid recruitment and organization of the actin cytoskeleton after adhesion. There, AGS cells appear fusiform, projecting cytoplasmic extensions indicating rearrangement similar to those induced by extracellular matrix proteins such as fibronectin (Fig. 6B). Cell spreading induced by CagI^C^ was significantly higher than CagI or CagL as seen by measurements of cell surface, cell perimeters or cell Feret’s diameter (Fig. S10). Next, we determined if K2 was able to inhibit CagI and CagI^C^ binding to AGS cells. After coating with the target proteins, the plates were incubated with increasing amount of K2 prior to incubation with AGS cells. As seen in Fig. 6C, DARPin K2 efficiently inhibits AGS cell binding to full-length CagI or CagI^C^ in a dose-dependent manner but had no effect on binding to CagL. In similar conditions, K11 showed a weak inhibition on cell adhesion on CagI. However, the inhibition was not dose-dependent, and no effect was observed on CagI^C^ (Fig. 6D). This suggests that the CagI epitopes targeted by K2 but not K11 are important for cell binding.

**Figure 6.**
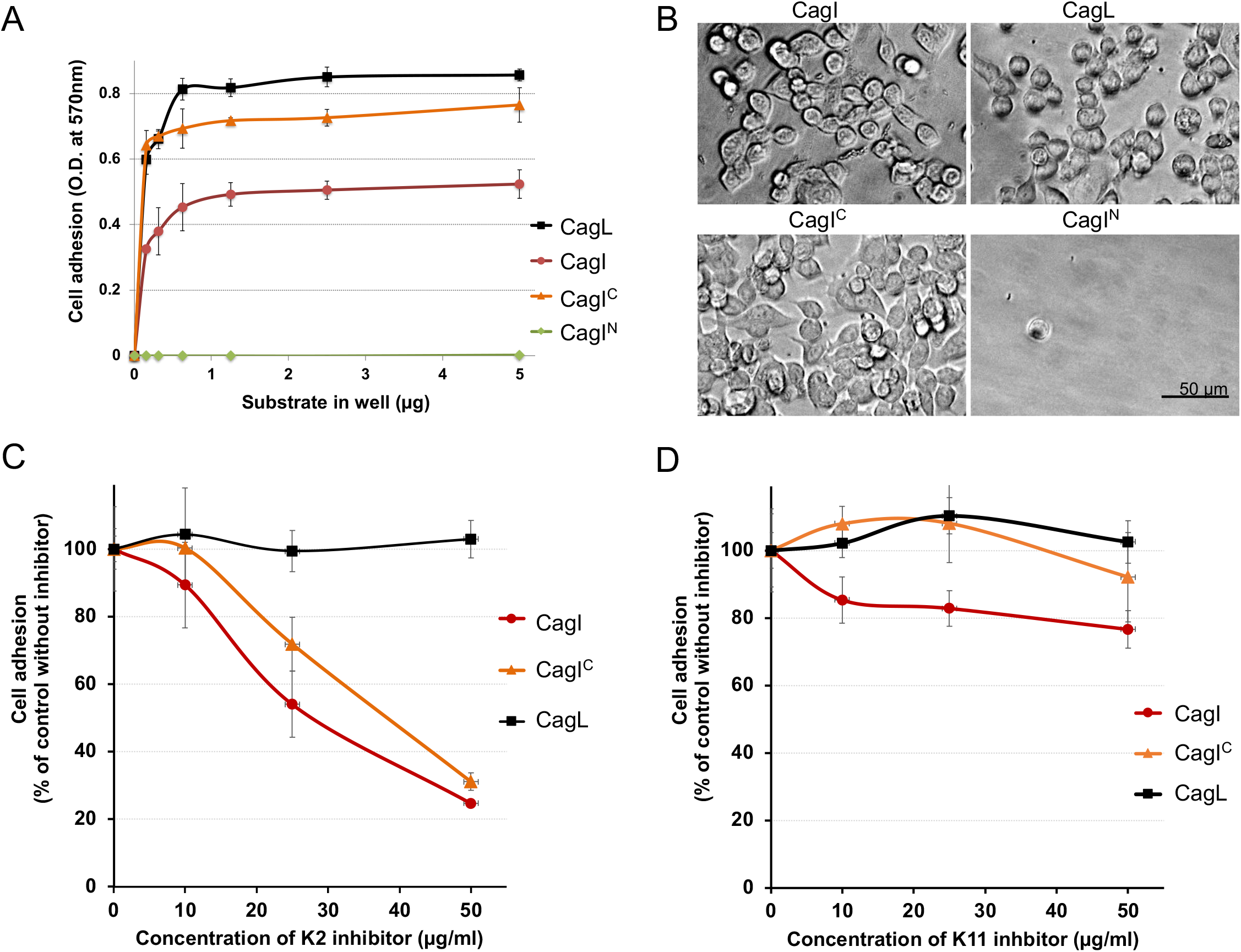
Cell binding and spreading to *cag*T4SS proteins and domains. A) Dose-dependent AGS cell adhesion to CagL, CagI, CagI^C^ and CagI^N^. Multiwell plates were coated with different amounts of the proteins as indicated on the Figure. Each assay point was derived from triplicate measurements. B) Representative images of adhered AGS cells on well-surfaces coated with 0.15 μg of indicated proteins. C) Effect of DARPin K2 or D) K11 on adhesion of AGS cells to CagL, CagI, CagI^C^. Multiwell plates were coated with 4 μg of each protein. After saturation with 1% BSA, the wells were incubated with 50 μL of the indicated concentration of K2 or K11 for 1 h at room temperature, and the cells were seeded in the presence of the inhibitor. The extent of adhesion was measured as previously described and expressed as percentage of adhesion to each protein in the absence of the inhibitor. Each assay point was derived from triplicate measurements.

## Discussion

*H. pylori cag*T4SS encodes for an unusually large number of proteins able to mediate host cell interaction, probably illustrating the long co-evolution of the bacteria with its host [39, 40]. Thanks to a remarkable genomic plasticity, the bacteria has evolved a plethora of mechanisms to adapt to human populations, and even different niches within an infected stomach [41]. While the CagA injection mechanism is still poorly understood, the *cag*T4SS pilus is essential to interact with the host cell and to deliver the oncoprotein [18]. Proteins associated with the *cag*T4SS pilus might have different functions. CagL not only interacts with integrins but also with the TLR5 receptor and this is also the case for CagY repeat region II [42, 43]. Structural and functional information is available on CagL [19, 24, 25], CagA [44, 45] and CagY [4, 11, 46], but little is known about the CagI protein despite its essential role in pilus formation [16].

Using an integrative structural biology approach, we show here that CagI forms an elongated dimer assembled via interactions between N-terminal domains followed by two C-terminal domains that are monomeric by themselves. On the one hand, the association mode of the N-terminal part was poorly predicted by AF, and although SAXS data confirms its general architecture, additional studies will be required to obtain structural details on this portion of CagI. On the other hand, our study, along with previous sequence analysis [16, 25], establish that CagL and the CagI C-terminal domain share structural similarity, including a set of three helices and a disulfide bridge. The resemblance between the two proteins extends to their localization: both were found present in the periplasm, and surface-exposed [30, 31]. CagL localizes at the tip of the pilus [21] but this is not entirely clear for CagI, although immunolocalization and electron microscopy identified CagI in *cag*T4SS pilus-like appendages [30]. Our work shows that similarly to CagL, CagI^C^ is able to mediate specific interactions with the host cell. Given that DARPin reducing this interaction also inhibit CagA injection, we assume that surface-exposed CagI interactions with host cell components are important for delivery through the *cag*T4SS.

Since we found that CagI and CagL interact *in vitro* [28] and *in vivo* [31] it is tempting to speculate that both CagL and CagI could be present at the surface or at the tip of the *cag*T4SS pilus where they could mediate adhesion and CagA translocation. Along these lines, both CagL and CagI were able to interact with integrin α_5_β_1_[28]. Here we show that CagI can mediate AGS cell attachment and spreading in a similar manner as CagL. However, the two proteins are unlikely to use the same determinant to interact with the host cell. CagL was proposed to interact with integrin via its RGD motif (reviewed in [18]) but this feature and surrounding motifs [18], are absent in CagI (Fig. S3). The CagL D1 motif involved in TLR5 recognition [43] is also not conserved in CagI (Fig. S3). Conversely, data presented in our study suggest that the motifs of CagI targeted by the K2 and K5 DARPins are important for host cell interactions, and these are also not conserved in CagL. The function of CagL and CagI are thus not redundant and instead might complement each other during *H. pylori cag*T4SS interaction with the host cell.

That CagI and CagL are surface associated proteins involved in host cell recognition is reminiscent of what is described in some other T4SSs. In *Agrobacterium tumefaciens*, VirB5 is located at the tip of the VirB/D T4SS pilus and but also promotes T-DNA translocation when added externally [47]. In the conjugative T4SSs from the plasmid pKM101, the VirB5 homologue TraC is also involved in host cell recognition and pilus adhesion [48]. TraC also interact with a protein named Pep (for PRD1 entry protein) with which it forms cluster at the surface of the bacterium to stimulate cell-to-cell contacts [49]. In the case of *H. pylori*, CagI and CagL, being exposed at the surface, might play a role in host-cell recognition. Indeed, while both proteins interact with integrins, the presence of these receptors is not essential for delivery of CagA from a mechanistic point of view [50]. However, *H. pylori* cells produce *cag*T4SS pilus and inject CagA at the basolateral surface, where integrins are located, but not at the apical sides of the cells [51]. Thus, CagI and CagL might sense the presence of integrins to trigger pilus assembly. Alternatively, CagI and CagL might be part of a larger, surface-accessible complex such as the lateral pores that have been observed on the *cag*T4SS pilus [14].

The results presented here identify a novel and effective target to prevent CagA delivery into host cell, hereby disarming the main oncogenic factor of *H. pylori*. Therapeutic strategies that target key virulence factors of pathogenic bacteria, while not actually killing the cells themselves, could prove to be vital for the treatment of numerous diseases [52]. In this regard, extracellular, species-specific appendages are appealing targets to prevent bacterial adhesion or effector translocation. Not only are they essential for virulence, but they have also the advantage of being easier to target, being accessible from the external milieu. Targeting pilus-associated proteins by small molecules or antibodies has been successful to disarm type III secretion systems [53]. In the case of *H. pylori cag*T4SS, efforts have so far been directed towards the screening or design of small molecules or peptides to target cytoplasmic components [54, 55]. Some inhibitors have shown some efficiency at inhibiting the cytoplasmic VirB11 ATPase [56, 57]. Other compounds inhibit pilus biogenesis and/or CagA translocation, but their mode of action remains undetermined [58]. The DARPin inhibition described here is a first example of effector translocation inhibition by a small protein binder targeting the external part of the *cag*T4SS. Although this inhibition is far from being complete, our study paves the way for strategies targeting *H. pylori*-specific extracellular determinants of CagA injection.

## Material and methods

### Protein structure prediction

We used the ColabFold Notebook for accessing AlphaFold2 Multimer [34, 59] to submit the CagI sequence (strain 26695 Genebank: AAD07606) to structure prediction for three models of a dimer. Other sequences were also used to evaluate the variability of CagI structures. Figures were generated with Pymol (Schrödinger) using the output PDB files containing the per residues LDDT scores.

### DNA manipulation, cloning, expression and protein purification

#### CagI, CagI_avi_ and CagI_strep_

Untagged CagI protein (residues 21 to 381) was purified as described in [28] using the pRSFMBP-*cagI* vector with the exception that detergent was removed by using an additional purification step. After HisMBP-tag cleavage CagI was loaded on a SOURCE™ 15Q 4.6/100 column (Cytiva) in a buffer containing 50 mM Tris pH 8, 20 mM NaCl and 0.004 % DDM and washed with buffer 50 mM Tris pH 8, 20 mM NaCl with 10 column volumes. The protein was eluted with a liner a gradient of buffer (50 mM Tris pH 8, 1M NaCl). The protein was then purified by size exclusion chromatography on a S200 Superdex column in 50 mM Tris pH 8, 150 mM NaCl.

The expression vector for CagI_avi_ was generated by insertion of an avitag (GLNDIFEAQKIEWHE) at the 3’ end of the *cagI* sequence in pRSFMBP-*cagI.* The CagI_avi_ protein was purified as above and biotinylated with the BirA enzyme as described in [60]. For CagI_Strep_ construction, expression and purification, the sequence corresponding to CagI residues 21-381 (from strain 26695) were fused to a C-terminal glycine-linker strep-tag encoding sequence (5’ GGTGGAGGTTCTGGCGGTGGATCGGGAGGTTCAGCGTGGTCTCATCCTCAATTTGAAAAA 3’). CagI_Strep_ was co-expressed with CagL_His_ protein (residues 21 to 237 fused to c-terminal his_6_-tag encoding sequence) in a pRSF duet vector. BL21 (DE3) cells carrying pRSF-CagI_Strep_-CagL_His_ expression vector were grown in LB at 37 °C until an OD_600_ of 0.6-0.8. Protein expression was induced for 16 h at 20 °C after adding 0.5 mM IPTG. For purification, cells were resuspended in lysis buffer (50 mM Tris pH 7.4, 200 mM NaCl) supplemented with protease inhibitor tablets (complete EDTA-free; Roche, one tablet per 250 mL of lysis buffer), lysozyme (0.1 mg/mL, Sigma-Aldrich) and DNase I (20 μg/mL, Sigma-Aldrich), and disrupted with three passages through a cell disrupter system (Constant Systems) operating at ~15,000 psi. The fraction containing CagI_Strep_ was separated from the soluble fraction after centrifugation at 7000 g for 20 min. The pellet was then resuspended in 50 mM Tris pH 7.4, 200 mM NaCl using a Dounce homogenizer and solubilized by addition of DDM to a final concentration of 0.5% by stirring at medium speed for 16 hours at 4°C. Insoluble material was pelleted by ultracentrifugation at 200,000 g for 1 hour at 4°C and the supernatant was loaded onto a StrepTrap column (Cytiva). The column was washed with 50 mM Tris pH 8, 150 mM NaCl, 0.005% DDM and CagI_Strep_ was eluted with 50 mM Tris pH 8, 150 mM NaCl, 0.005% DDM supplemented with 2.5 mM desthiobiotin. The elution fraction was analysed by SDS-PAGE before to loading on a HisTrap column (Cytiva) to remove CagL_His_ proteins. The flow-through, containing CagI_Strep_ was collected, concentrated and loaded onto a Superdex 200 HiLoad 16/600 gel filtration column (Cytiva) equilibrated in 50 mM Tris pH 8, 150 mM NaCl and 0.005 % DDM.

#### CagI domains

PCR fragments encoding for CagI residues 27-190 (CagI^N^) or residues 191 to 381 (CagI^C^) were amplified using primers *cagI^N^fw* (5’-CACCACGCTTGAACCCGCCTTAAAAG-3’), *cagI^N^rev* (5’- TCAACTTCCTAGAGCTTGAGAAAG), *cagI^C^fw* (CACCTCTTCTGACAACGCTCAATACATC), *cagI^C^rev* (TCATTTGACAATAACTTTAGAGCTAG) and inserted into the pET151D topo vector (ThermoScientific) following the manufacturer’s protocol. The resulting vectors pET151CagI^N^ or pET151CagI^C^ encode each CagI domain fused to a N-terminal His_6_-tag followed by a tobacco-etch virus (TEV) cleavage site. *E. coli* T7 Express cells harboring pET151CagI^N^ or pET151CagI^C^ were grown in 1 L LB medium supplemented with ampicillin (100 μg/ml) at 37 °C until OD_600nm_ reached 0.6. Protein expression was induced for 16 hours at 20° C by adding 1 mM IPTG (final concentration). Cells were harvested and resuspended in 20 ml of buffer AG (50 mM Tris pH 8, 200 mM NaCl, 5 % glycerol (v/v)). Solutions containing the cells were supplemented with Triton-X100 (final concentration 1 %), one tablet (per 250 ml of buffer) of complete EDTA-free protease inhibitor (Roche), lysozyme (0.1 mg/ml, Sigma-Aldrich) and DNase I (20 μg/ml, Sigma-Aldrich) prior to sonication. Cell debris was removed by centrifugation (14,000 g, 4° C, 15 min) and the supernatants were loaded onto a HisTrap column (Cytiva). Proteins were eluted with a 0 to 100 % linear gradient of buffer AG containing 500 mM imidazole. Fractions containing CagI domains were pooled and His_6_ tags were cleaved by TEV protease with 0.5 mM EDTA and 5 mM DTT and dialysed 16 h against buffer AG at 4°C. Proteins were loaded onto HisTrap column and flowthrough containing cleaved protein was pooled and concentrated (Amicon 3 K Sigma Aldrich). Proteins were loaded onto a Superdex 200 increase 10/300 GL column (Cytiva) equilibrated in buffer AG.

### Selection and screening of DARPins

To generate DARPin binders, CagI biotinylated at a C-terminal avi tag was immobilized, in alternating selection rounds, on either MyOne T1 streptavidin-coated beads (Pierce) or Sera-Mag neutravidin-coated beads (GE). Ribosome display selections were performed essentially as described [37], using a semi-automatic KingFisher Flex MTP96 well platform.

The library includes N3C-DARPins with the original randomization strategy as reported [61] but also a stabilized C-cap [33, 62, 63]. Additionally, the library is a mixture of DARPins with randomized and non-randomized N- and C- terminal caps, respectively [33, 64]. Successively enriched pools were ligated as intermediates in a ribosome display-specific vector [64]. Selections were performed over four rounds with decreasing target concentration and increasing washing steps, and the third round included a competition with non-biotinylated CagI to enrich for binders with high affinities.

The final enriched pool of cDNA encoding putative DARPin binders was cloned as fusion construct into a bacterial pQE30 derivative vector (Qiagen), containing a T5 lac promoter and lacIq for expression control, with a N-terminal MRGS(H)_8_ tag and C-terminal FLAG tag via unique *Bam*HI and *Hind*III sites. After transformation of *E. coli* XL1-blue, 380 single DARPin clones selected to bind CagI were expressed in 96-well format and lysed by addition of B-Per Direct detergent plus lysozyme and nuclease (Pierce). After centrifugation these crude extracts were used for initial screening to bind CagI using ELISA. For IMAC purification of DARPins they were expressed in deep-well 96-well plates, lysed with Cell-Lytic B (SIGMA) and purified over a 96-well IMAC column (HisPur™ Cobalt plates, Thermo Scientific).

ELISAs were performed using streptactin-coated plates (iba-lifesciences) and used for immobilization of CagI_strep_ at a concentration of 50 nM. Detection of DARPins (1:1000 dilution of crude extracts, or a concentration of 50 nM for IMAC-purified DARPins) binding to CagI was performed using a mouse-anti-FLAG M2 monoclonal antibody (dilution 1:5000; Sigma, F1804) as primary and a goat-anti-mouse antibody conjugated to an alkaline phosphatase (dilution 1:10,000; Sigma, A3562) as secondary antibody. After addition of pNPP (para-nitrophenyl phosphate) absorbance at 405 nm was determined after 30 minutes. Signals at 540 nm were subtracted as background correction.

### DARPin purification and pull downs assays

pQE30 expression vectors expressing the DARPins K5, K9, K2, K12, K15, K8, K10 and K11 (described above) were transformed into T7 Express cells. Protein expression and extraction were performed as described above for CagI domains in buffer A (50 mM Tris pH 8, 150 mM NaCl) for K5 or buffer AG (50 mM Tris pH 8, 150 mM NaCl, glycerol 5 % (v/v)) for others DARPins. Supernatants from 1L bacterial cell cultures were loaded onto a HisTrap column (Cytiva), washed successively with 3 column volumes of buffer AG (or buffer A for K5) 2 column volumes of buffer AG (or A for K5) supplemented with 1 M NaCl. Proteins were eluted with a 0 to 100 % linear gradient of buffer AG (or buffer A for K5) containing 500 mM imidazole. Fractions containing the DARPins were pooled, concentrated (Amicon 3 K Sigma Aldrich) and loaded onto a Superdex 200 increase 10/300 GL column (GE healthcare) equilibrated in buffer AG or buffer A (for K5).

CagI_Strep,_ CagI^N^ and CagI^C^ were mixed and incubated on ice for 1 h with 50 μg of _His8_DARPins and 2-fold molar excess of CagI. The mixtures were incubated with 30 μL Ni-NTA magnetic beads (Merck) and loaded onto Biosprint 15 (Qiagen) for purification. Proteins were washed twice with 750 μL of buffer AG or A (for CagI:K5) and finally eluted in 150 μL buffer A (CagI:K5) or AG containing 500 mM imidazole.

### Co-expression of DARPins and CagI_strep_

pRSF-*cagI_strep_* and pQE30-DARPin vectors were introduced in *E. coli* T7 Express cells (NEB). Cells were grown in 50 mL LB medium supplemented with kanamycin (50 μg/mL) and ampicillin (100 μg/mL) at 37 °C until OD_600nm_ reached 0.6. Protein expression was induced for 16 hours at 20° C by adding 1 mM IPTG (final concentration). Cells were harvested and resuspended in 1 ml of buffer A (50 mM Tris pH 8, 150 mM NaCl) for CagI: K5 or buffer AG for the remaining CagI:DARPin complexes. For cell lysis buffers were supplemented with Triton-X100 (final concentration 1%), one tablet (per 250 mL of buffer) of complete EDTA-free protease inhibitor (Roche), lysozyme (0.1 mg/mL, Sigma-Aldrich) and DNase I (20 μg/mL, Sigma-Aldrich). The cells were sonicated and centrifuged at 14,000 g 4° C for 15 min. Supernatants were incubated with 30 μL nickel magnetic beads (Merck) and loaded onto Biosprint 15 (Qiagen). Proteins were washed two times with 750 μl of buffer A (or AG) and eluted with 150 μL buffer A (or AG) supplemented with 500 mM imidazole. For western-blot detection, proteins samples were separated on a 20% SDS-PAGE, transferred to a polyvinylidene difluoride membrane, and immunoblotted using a mouse monoclonal antibody against Strep-tag (Qiagen, 34850). Alkaline phosphatase conjugated to anti-mouse IgG (Sigma-Aldrich, A1293) was used as a secondary antibody. Detection was performed by colorimetry using nitro blue tetrazolium chloride / 5-bromo-4-chloro-3-indolyl-phosphate (NBT-BCIP, Sigma-Aldrich) as a substrate.

For large scale purification, CagI:DARPins expression and extraction of proteins were performed as described above except for a larger volume of cell culture (1 L). After lysis by sonication and centrifugation (14,000 g, 4°C 15 minutes), the CagI:DARPins supernatants were loaded onto a 5 mL StrepTrap column (Cytiva), washed with 3 column volumes of buffer AG (or A for CagI:K5) and eluted with buffer AG (or buffer A for the CagI:K5 complex) containing 2.5 mM desthiobiotin. Elution fractions were loaded onto a 5 mL HisTrap column (Cytiva) and after a 3 column volumes wash with buffer AG (or A for CagI:K5), proteins were eluted in buffer AG (or A, see above) containing 500 mM imidazole. Fractions containing the complexes were concentrated (Amicon 10 K Sigma Aldrich) and loaded onto a Superdex 200 increase 10/300 GL column (Cytiva) equilibrated in 50 mM Tris pH 8, 150 mM NaCl for CagI:K5 complex and 50 mM Tris pH 8, 200 mM NaCl, 5 % glycerol v/v for other CagI:DARPins complexes.

### Multi-angle light scattering (MALS)

Size-exclusion chromatography experiments coupled to multi-angle laser light scattering (MALS) and refractometry (RI) were performed on a Superdex S200 Increase 5/150 GL column with size-exclusion buffer 50 mM Tris pH 8, 150 mM NaCl for DARPin K5 and CagI-K5 complex and with buffer 50 mM Tris pH 8, 200 mM NaCl, 5% glycerol v/v for others CagI-DARPin complexes. Fifty microliters of proteins were injected at a concentration of 5 to 8 mg/mL. Online MALS detection was performed with a miniDAWN-TREOS detector (Wyatt Technology Corp., Santa Barbara, CA, USA) using a laser emitting at 690 nm and by refractive index measurement using an Optilab T-rEX system (Wyatt Technology Corp.). Weight-averaged molar masses (Mw) were calculated using the ASTRA software (Wyatt Technology Corp.).

### Crystallization, structure determination and refinement

Crystals of CagI:K5 and CagI:K2 were obtained by the sitting drop vapor diffusion method using a Mosquito robot. Drops consisting of 200 nL of protein complex (7 mg/mL) with 200 nL of reservoir solution were left at 19°C for two weeks. Crystals of CagI:K5 appeared in condition E5 of the PACT Premier™ screen (Molecular Dimension) with a reservoir solution consisting of 0.2 M sodium nitrate, 20 % w/v PEG 3350). CagI:K2 crystals appeared in condition E2 of the same screen with a reservoir solution consisting of 0.2 M sodium bromide, 20 % w/v PEG 3350). Crystals were flash frozen in reservoir solution supplemented with glycerol 15% (v/v). Data were collected at 100 °K at PROXIMA 1 beamline of the synchrotron SOLEIL and processed using XDS [65] and AIMLESS [66] from the CCP4 program suite [67, 68]. Crystals of CagI:K5 and CagI:K2 diffracted to resolutions of 2.0 Å and 1.8 Å, respectively and belonged to the orthorhombic space group P2_1_2_1_2_1_ with very similar cell dimensions (Table 2). The structure of CagI:K5 was solved by molecular replacement using the coordinates of DARPin E11 (PDB ID: 6FP8 [69]) as a probe in PHASER [70]. Examination of the resulting electron density indicated that additional helices were present in the asymmetric unit. Manual building resulted in a first model of several helices and the unit contained one K5 molecule and a fragment of CagI. After several rounds of manual building/refinement the sequence of the additional peptide could be undoubtedly attributed to CagI residues 204 to 307. The resulting model was used as a template to solve the structure of CagI:K2. The models were refined with final R_work_/R_free_ of 0.18/0.23 (CagI:K5) and 0.17/0.20 (CagI:K2). The coordinates and structure factors were deposited in the Protein Data Bank with accession code 8AIW (CagI:K5) and 8AK1 (CagI:K2).

### Small-angle X-ray scattering

SAXS data were collected for CagI and CagI domains at the ESRF BioSAXS beamline BM29 using an online size-exclusion chromatography setup. Fifty μl of protein (8 mg/mL) were injected into a size-exclusion column (S200 increase 5/150) equilibrated in 50 mM Tris, pH 8.0, 200 mM NaCl, 5% glycerol v/v. Images were acquired every second for the duration of the size-exclusion run. Buffer subtraction was performed by averaging 100 frames. Data reduction and analysis was performed using the BsxCuBE data collection software and the ATSAS package [71]. The program AutoGNOM was used to generate the pair distribution function (*P*(*r*)) and to determine *D*_max_ and *R_g_* from the scattering curves (*I*(*q*) *versus q*) in an automatic, unbiased manner. Theoretical curves from the models were generated by FoXS [72]. Ab initio modelling was performed with DAMMIN [73].

### Surface plasmon resonance

Measurements were performed using a Biacore T200 instrument (Cytiva). CagI and CagI^C^ were covalently immobilised to the dextran matrix of a CM5 sensorchip via their primary amine groups. The carboxymethylated dextran surface was activated by the injection at 5 μL/min of a mixture of 200 mM EDC [*N*-ethyl-*N*′-(3-dimethylaminopropyl)carbodiimide] and 50 mM NHS (*N*-hydroxysuccinimide). Ligands were diluted in 10 mM sodium acetate pH 4 to a 10-20 μg/mL concentration before injection over the activated surface of the sensor chip. Residual active groups were blocked by injection of 1 M ethanolamine pH 8.5. Immobilization levels of 1,200 RU (response units) were obtained for CagI and 340 RU for CagI^C^. A control flow cell was activated by the NHS/EDC mixture and deactivated by 1 M ethanolamine pH 8.5 without any coupled protein. Control sensorgrams were subtracted online from the sensorgrams to derive specific binding responses. Analytes were injected at 50 μg/mL for 120 seconds after dilution in running buffer (10 mM HEPES pH 7.4, 150 mM NaCl, 0.05 % P20). The sensorchip surface was regenerated with 2 pulses (30 sec) of ethylene glycol 50 % and 1 pulse of 2 M guanidine hydrochloride. The equilibrium K_D_ were calculated using the Biacore T200 evaluation software (v3.2.1).

### Quantification of CagA translocation and bacterial cell binding

Translocation of CagA into AGS cells was determined quantitatively using either the TEM-1-CagA translocation assay [37], or the HiBiT-CagA translocation assay [38]. For the TEM-1-CagA assay, AGS cells were co-incubated with *H. pylori* P12 [TEM-1-CagA] for 2.5 h in 96-well microtiter plates in PBS/10% FCS. After infection, cells were loaded with fluorescent substrate CCF4-AM in a loading solution (LiveBLAzer-FRET B/G loading kit; Invitrogen) supplemented with 1 mM probenecid (Sigma) according to the manufacturer’s instructions. Cells were incubated with this loading solution at room temperature in the dark for 2 h, and then measured with a Clariostar reader (BMG Labtech) using an excitation wavelength of 405 nm, and emission wavelengths of 460 nm, or 530 nm. CagA translocation was calculated as the ratio of background-corrected emission values at 460 nm to 530 nm and normalized to *H. pylori* P12 [TEM-1-CagA] and P12Δ*cagT* [TEM-1-CagA] as positive and negative controls. For the HiBiT-CagA assay, *H. pylori* P12 [HiBiT-CagA] was pre-cultured in PBS/10 % FCS for 2 h at 37 °C, 10% CO_2_. Subsequently, AGS [LgBiT] cells seeded in a 96-well plate (4titude) were infected with 200 μl of this pre-culture, and incubated at 37 °C, 5 % CO_2_ for 2.5 h. Supernatants containing unbound bacteria were discarded, and cells were loaded with 40 μL PBS/FCS and 10 μL 5x luciferase substrate mix. After 10 min incubation, luminescence was measured at 470 nm in a Clariostar reader. The amount of translocated HiBiT-CagA was calculated after correction for the background signal as percentage in relation to the untreated *H. pylori* P12 [HiBiT-CagA] control. For inhibition experiments, bacteria were pre-incubated with the respective DARPins at a concentration of 5 μM for 30 minutes at 37 °C in PBS/10% FCS, followed by infection for 2.5 h in the presence of the DARPins.

For determination of bacterial binding to gastric cells, AGS cells were infected with *H. pylori* strain P12 [pHel12∷gfp] [74], using an MOI of 60, and infection was allowed to proceed for 1 h at 37 °C and 5% CO_2_. For inhibition experiments, bacteria were pre-incubated with the respective DARPins at a concentration of 5 μM for 30 minutes at 37 °C in PBS/10% FCS, and then co-incubated with AGS cells, as above. After three washing steps with PBS, cells with adherent bacteria were collected by EDTA treatment, and analysed for GFP fluorescence in a flow cytometer (FACS CantoII, BD Biosciences). For analysis, the median fluorescence intensity of non-infected cells was subtracted from that of infected samples.

### Cell Adhesion and Inhibition Assays

Gastric adenocarcinoma cell line AGS (CRL 1739; American Type Culture Collection) was grown in F-12K Medium (Kaighn’s Modification of Ham’s F-12 Medium) supplemented with 10% fetal calf serum. Tissue culture 96-well plates (Nunc PolySorp, Dutscher, France) were coated with serial dilutions of the indicated proteins, CagL, CagI, CagI^C^ and CagI^N^ (0-5 μg/well in PBS) by overnight adsorption at 4°C. The amount of adsorbed protein was determined with a BCA microprotein assay. After saturation of the wells with 1% BSA, AGS cells were collected from the culture plates by detaching with 5 mM EDTA/PBS, followed by rinsing and suspending in F-12K serum-free medium. AGS cells were seeded on ligand-coated plates at 8 × 10^4^ cells/well. After 1 to 2 h, nonadherent cells were removed with a PBS wash. The extent of adhesion was determined by fixing adherent cells with 1% glutaraldehyde in PBS and then staining with 0.1% crystal violet and measuring absorbance at 570 nm as described previously [75, 76]. A blank value was subtracted that corresponded to BSA-coated wells. Each assay point was derived from triplicate measurements (three wells per assay point). Adherent cells were photographed 1 hour after seeding with an Axiovert 40 Zeiss microscope equipped with Differential Interference Contrast coupled to a Coolsnap Fx Camera (Roper Scientific, Evry, France). Cell size measurements were performed manually using Fiji 1.53c (plus) software. For AGS cell adhesion inhibition experiments with inhibitors, the coated wells were incubated with serial dilutions in PBS for 60 min at room temperature prior to the adhesion assay as indicated in the corresponding figures. Cell adhesion data are presented as the means ± SD using Excel. For cell size measurements, a one-way Anova test was used for groups comparisons using Prism (GraphPad) software. The significance threshold was set for the t-test as P <0.05. The exact sample size for replicate measurements is specified in each graph legend.

## Acknowledgments

This project was funded by the ANR-Subsist (ANR- 19-CE11-0012) and the Ligue Regionale Isère Contre le Cancer. We acknowledge the contribution of the Protein Science Facility of the SFR Biosciences (UMS3444/US8). We thank Thomas Reinberg and Joana Marinho for expertly carrying out many steps in the DARPin selection. We thank Dr. Petya Pernot from the BM29 beamline at the synchrotron ESRF for help with SAXS data collection and processing. Thanks are also due to PROXIMA 1 beamline staff for help in data collection and processing.

## Authors contribution

MB, JG, AV, CL, RF, SF, BD, CB, PR, LT: Investigation

AP, WF, LT , PR, SVLG: Formal Analysis, Conceptualization.

AP, RF, WF, LT, PR: Funding acquisition

LT, MB: Writing – Original Draft Preparation

WF, PR, SVLG, LT, AP: Writing – Review & Editing

