## Supplementary information for "Designed Ankyrin Repeat Proteins provide insights into the structure and function of CagI and are potent inhibitors of CagA translocation by the *Helicobacter pylori* type IV secretion system"

**Contains**

- Tables S1
- Supplementary Figures legends
- Figures S1-S10

**Table S1. Small angle X-ray scattering data collection, processing and analysis**

| | CagI | $CagI^N$ | $CagI^C$ |
|---|---|---|---|
| **Data-collection parameters:** | | | |
| Q range ($Å^{-1}$) | 0.01-0.17 | 0.01-0.18 | 0.01-0.23 |
| **Structural parameters:** | | | |
| I(0) ($Å^{-1}$) (from P(r)) | 18.2 | 5.7 | 9.4 |
| Rg (Å) (from P(r)) | 4.66 ± 0.03 | 2.9 ±0.1 | 2.4 ± 0.02 |
| I(0) ($Å^{-1}$) (from Guinier) | 18.2 | 5.7 | 9.4 |
| Rg (Å) (from Guinier) | 4.65 ± 0.07 | 2.9 ±0.1 | 2.3 ± 0.02 |
| Dmax (Å) | 162 | 116 | 88 |
| Porod volume estimation. $V_0$ ($Å^3$) | 155.3 | 54.4 | 27.8 |
| **Molecular-mass determination:** | | | |
| Molecular mass Mr (kDa) (SaxsMOW) | 99.1 | 32 | 20.3 |
| Molecular mass Mr (kDa) (Porod) | 91.3 | 32 | 16.3 |
| Calculated mass Mr (kDa) (sequence) | 78.2 | 35.6 | 18.1 |

**Supplementary Figure legends**

**Figure S1. Structural modeling of CagI dimer.** Scores and sequence coverage of the AF models (left) and cartoon depiction of the AF models of CagI dimer (right) coloured according to pLDDT scores (30 to 100).

**Figure S2. Small Angle X-ray Scattering study of CagI, CagI$^N$ and CagI$^C$.** Size exclusion coupled to experimental SAXS curves of CagI (black), CagI$^N$ and CagI$^C$ compared to theoretical curves obtained with the corresponding model 1, 2 and 3 and domains. Theoretical curves were obtained with CagI dimers, CagI$^N$ dimers and CagI$^C$ monomers (chain A). Below are cartoon representations of the structures of model 1 (orange), model 2 (blue) and model 3 (magenta) fitted into the DAMMIN envelope obtained for each SAXS data. From left to right: full length CagI dimer, CagI$^N$ dimer and CagI$^C$ monomer.

**Figure S3. Sequence conservation of CagI.** Clustal O alignment of CagI sequences. CagI sequence from strain 26695 (Uniprot O25273) is labelled CagI. The other CagI sequences are labelled with their uniprot code and have been chosen to illustrate the low diversity. CagL sequence corresponds to the one from strain 26695 (Uniprot O25272). Secondary structures of CagI (AF model) and CagL (PDB code 3ZCJ) are indicated above and below the alignment, respectively. The predicted signal peptide sequence (SP) is indicated by a blue box, the cysteines forming disulfide bridges are indicated by a blue dot. The arginine-glycine-aspartate [21] and D1 motifs [43] of CagL are indicated by magenta and green lines, respectively.

**Figure S4. Validation of binding of $_{His8}$-DARPins to CagI$_{Strep}$ by co-expression and co-purification.** Coomassie blue stained SDS-PAGE of A) *E. coli* cell extracts before (-), after (+) induction of CagI$_{strep}$ expression and (E) elution fraction of His-trap column. No CagI protein could be detected in this fraction but a contaminant of around 39 kDa is visible and indicated by a *. B) co-purification of $_{His8}$-DARPins (K1-K15) with CagI-$_{Strep}$ on Ni-NTA beads showing that a band corresponding to CagI$_{strep}$ is present in the elution fraction when co-expressed with K2, K5, K9, K12, K8, K10, K11 and K15 but not with K3, K4, K6, K7, K13, K14 and K1. C) Western-blot analysis of the same samples using anti-strep antibody and stained with NBT-BCIP. The band corresponding to CagI$_{strep}$ is indicated by an arrow.

**Figure S5. Sequences of DARPins identified in this study.** Sequences of the DARPins, with name and clone ID indicated, aligned using ClustalO. Amino-acid differences are shaded in grey except for the single difference between K2 and K5 shaded in cyan. Repeat, N-cap and C-cap regions are indicated above the sequences.

**Figure S6. Affinity of DARPin for CagI and CagI$^{C}$.** Surface Plasmon Resonance experiments using single-cycle-mode on CM5 chips coated with CagI (A) or with (B) CagI$^{C}$. DARPins were injected on the chips at increasing concentrations as follows. For experiments performed on full-length CagI, concentrations of DARPins were: 0.5, 2.5, 12.5, 62.5 and 312.5 nM for K9, K11, K12, K15; and 1, 3, 9, 27 and 81 nM for K10. For CagI$^{C}$ experiments, injections of DARPins were performed with concentrations of 0.05, 0.15, 0.45, 1.35 and 4 nM for K9, K10 and K11; 0.16, 0.8, 4, 20 and 100 nM for K12; and 2, 4, 8, 16 and 32 nM for K15. Fit curves obtained with heterogenous ligand model (CagI) or binding model 1:1 (CagI$^{C}$) are shown as black dashed lines.

**Figure S7. Structural comparison of crystal structure of CagI$^{204\text{-}307}$ and corresponding region in the AF model.** Two rotated views of the crystal structure of CagI$^{204\text{-}307}$ from the CagI:K2 complex (green) and AF model (grey) displayed as cartoon. The side chains of the cysteine residues 272 and 283 forming the disulfide bond are displayed as ball-and-stick with sulfur atoms coloured in yellow.

**Figure S8. Determination of molar mass of CagI:DARPins complexes by size exclusion chromatography coupled to multi-angle light scattering.** Each purified CagI:DARPin complex was submitted to SEC-MALS measurement represented by A280 chromatograms. Molar mass calculations are represented by dotted lines on each graph.

**Figure S9. Cell binding measurements by flow cytometry with *H. pylori* P12 [pHel12::gfp]** pre-treated with DARPins or left untreated. Data shown are mean values and standard deviations of median fluorescence intensities (MFI) normalized to untreated bacteria, resulting from three independent experiments.

**Figure S10. Quantitative measurement of cell spreading on CagL, CagI or CagI$^{C}$.** Morphological characterization of cells adhered to 0.15 µg of CagL, CagI, CagI$^{C}$ 60 minutes after seeding. Cell surface area, perimeter and Feret's diameter were determined on phase contrast images using Fiji software. Each dot represents one cell (60 adhered AGS cells were measured for each condition). Each of the three parameters clearly shows that cells that have adhered to CagI$^{C}$ have a larger and more extended contact surface compared to cells bound to CagI or CagL. A one-way ANOVA with Tukey's post-test was used to determine significance. Means ± SD are shown with *p<0.1, ****p<0.0001.

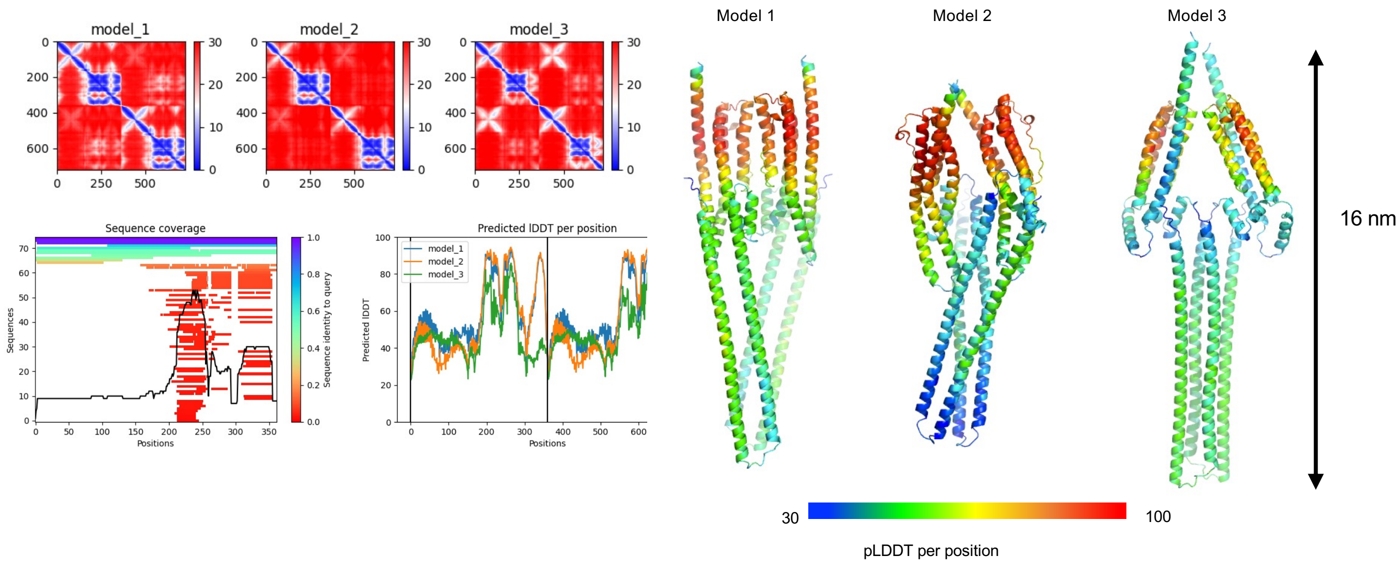


**Figure S1. Structural modeling of CagI dimer.** Scores and sequence coverage of the AF models (left) and cartoon depiction of the AF models of CagI dimer (right) coloured according to pLDDT scores (30 to 100).

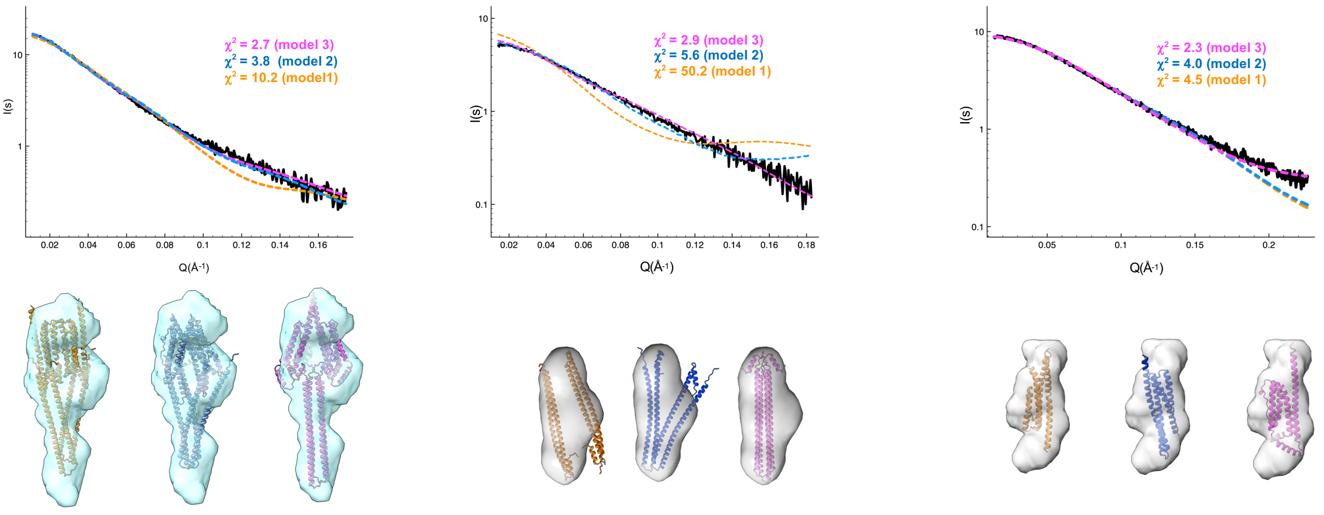


CagI (dimer) | CagI$^{N}$ (dimer) | CagI$^{c}$ (monomer)

**Figure S2. Small Angle X-ray Scattering study of CagI, CagI$^{N}$ and CagI$^{C}$.** Size exclusion coupled to experimental SAXS curves of CagI (black), CagI$^{N}$ and CagI$^{C}$ compared to theoretical curves obtained with the corresponding model 1, 2 and 3 and domains. Theoretical curves were obtained with CagI dimers, CagI$^{N}$ dimers and CagI$^{C}$ monomers (chain A). Below are cartoon representations of the structures of model 1 (orange), model 2 (blue) and model 3 (magenta) fitted into the DAMMIN envelope obtained for each SAXS data. From left to right: full length CagI dimer, CagI$^{N}$ dimer and CagI$^{C}$ monomer.

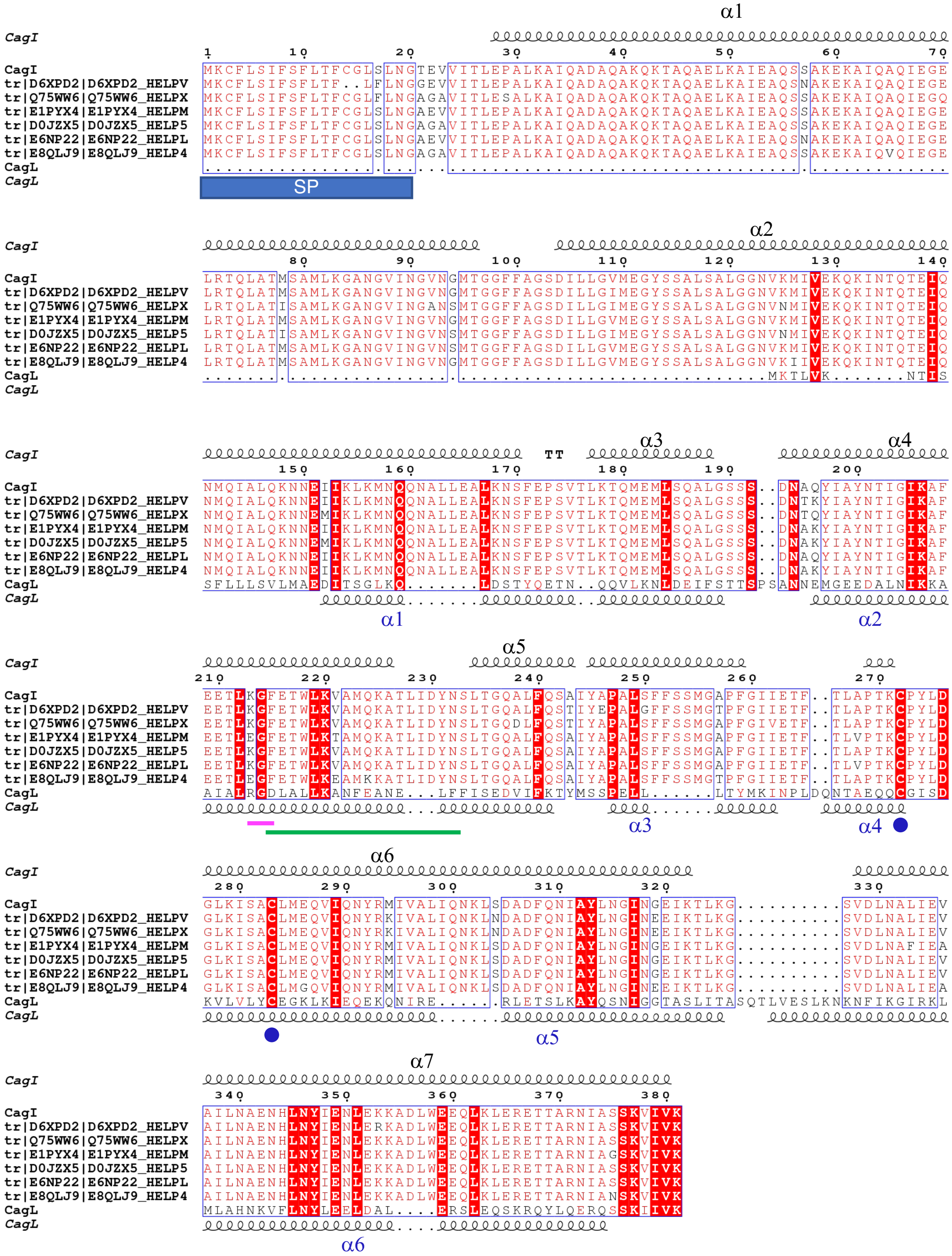


**Figure S3. Sequence conservation of CagI.** Clustal O alignment of CagI sequences. CagI sequence from strain 26695 (Uniprot O25273) is labelled CagI. The other CagI sequences are labelled with their uniprot code and have been chosen to illustrate the low diversity. CagL sequence corresponds to the one from strain 26695 (Uniprot O25272). Secondary structures of CagI (AF model) and CagL (PDB code 3ZCJ) are indicated above and below the alignment, respectively. The predicted signal peptide sequence (SP) is indicated by a blue box, the cysteines forming disulfide bridges are indicated by a blue dot. The arginine-glycine-aspartate [21] and D1 motifs [43] of CagL are indicated by magenta and green lines, respectively.

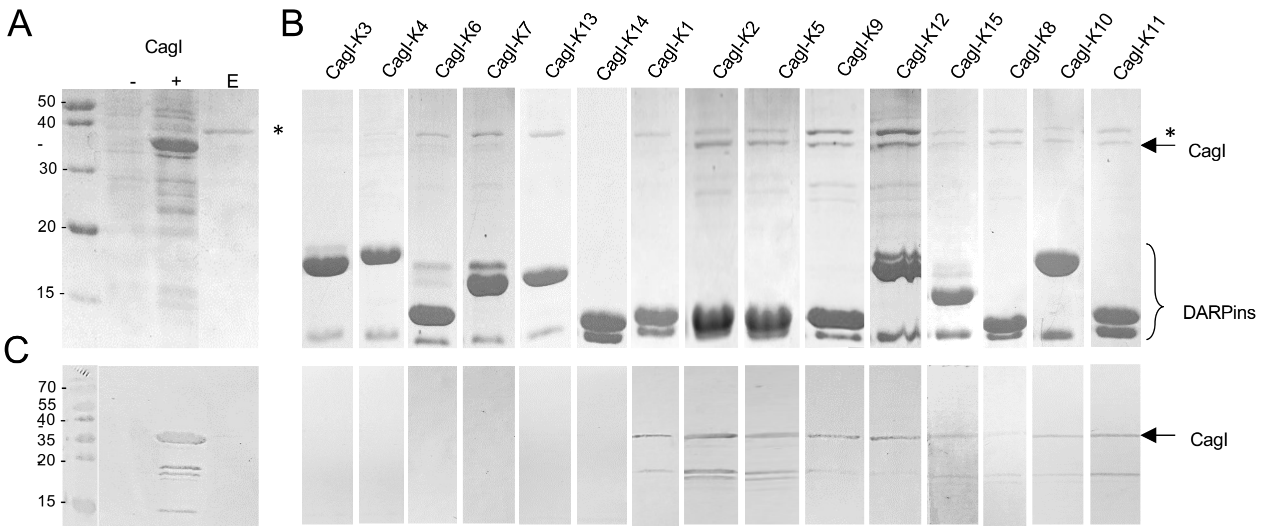


**Figure S4. Validation of binding of $_{His8}$-DARPins to $CagI_{Strep}$ by co-expression and co-purification.** Coomassie blue stained SDS-PAGE of A) *E. coli* cell extracts before (-), after (+) induction of $CagI_{strep}$ expression and (E) elution fraction of His-trap column. No CagI protein could be detected in this fraction but a contaminant of around 39 kDa is visible and indicated by a *. B) co-purification of $_{His8}$-DARPins (K1-K15) with $CagI\text{-}_{Strep}$ on Ni-NTA beads showing that a band corresponding to $CagI_{strep}$ is present in the elution fraction when co-expressed with K2, K5, K9, K12, K8, K10, K11 and K15 but not with K3, K4, K6, K7, K13, K14 and K1. C) Western-blot analysis of the same samples using anti-strep antibody and stained with NBT-BCIP. The band corresponding to $CagI_{strep}$ is indicated by an arrow.

| DARPin | Clone ID | N-cap | 1st repeat |
|---|---|---|---|
| K2 | 008-801-2176-F2 | DLGKKLLEAA LIGQDDEVRI LMANGADVNA | MDNFGHTPLH LAAMMGHLEI VEVLLKTGAD VNA |
| K5 | 008-801-2177-B8 | DLGKKLLEAA LIGQDDEVRI LMANGADVNA | MDNFGHTPLH LAAMMGHLEI VEVLLKTGAD VNA |
| K8 | 008-801-2177-G11 | DLGKKLLEAA TIGQHDEVRI LMANGADVNA | WDFLGQTPLH LAANMGHLEI VEVLLKAGAD VNA |
| K9 | 008-801-2178-B4 | DLGKKLLEAA RAGQDDEVRI LMANGADVNA | TDMAGWTPLH LAAIEGHLEI VEVLLKTGAD VNA |
| K10 | 008-801-2178-C11 | DLGKKLLEAA VQGQDDEVRI LMANGADVNA | QDQAGHTPLH LAAVRGHLEI VEVLLKTGAD VNA |
| K11 | 008-801-2178-E11 | DLGKKLLEAA LYGQDDEVRI LMANGADVNA | QDQAGHTPLH LAALVGHLEI VEVLLKTGAD VNA |
| K12 | 008-801-2179-E4 | DLGKKLLEAA HAGQDDEVRI LMANGADVNA | SDIWGFTPLH LAALWGHLEI VEVLLKTGAD VNA |
| K15 | 008-801-2179-E12 | DLGKKLLEAA SWGQDDEVRI LMANGADVE- | ----GYTPLH LAASWGHLEI VEVLLKTGAD VNA |

| DARPin | Clone ID | 2nd repeat | 3rd repeat |
|---|---|---|---|
| K2 | 008-801-2176-F2 | FDLTGFT PLHLAAYAGH LEIVEVLLKH GADVNA---- | ---------- ---------- ----------- |
| K5 | 008-801-2177-B8 | FDLTGFT PLHLAAYAGH LEIVEVLLKH GADVNA---- | ---------- ---------- ----------- |
| K8 | 008-801-2177-G11 | EDNHGFT PLHLAAYWGH LEIVEVLLKH GADVNA---- | ---------- ---------- ----------- |
| K9 | 008-801-2178-B4 | EDAFGLT PLHLAAWFGH LEIVEVLLKH GADVNA---- | ---------- ---------- ----------- |
| K10 | 008-801-2178-C11 | TDDAGWT PLHLAAMHGH LEIVEVLLKA GADVNATDVW | GHTPLHLVAV REGHLEIVEV LLKHGA DVNA |
| K11 | 008-801-2178-E11 | ADDFGFT PLHLAAFYGH LEIVEVLLKH GADVNA---- | ---------- ---------- ----------- |
| K12 | 008-801-2179-E4 | NDSFGET PLHLAASFGH LEIVEVLLKA GADVNAIDYF | GWTPLHLAAV -NGHLEIVEV LLKTGA DVNA |
| K15 | 008-801-2179-E12 | HDMNGFT PLHLAAFYGH LEIVEVLLNA GADVNAQDYQ | GNTPLHLAAM -FGHLEIVEV LLKHGA DVNA |

| DARPin | Clone ID | C-cap |
|---|---|---|
| K2 | 008-801-2176-F2 | QDQDGATPFD LAAWFGNEDI AEVLQKAAKL |
| K5 | 008-801-2177-B8 | QDQDGATPFD LAAWLGNEDI AEVLQKAAKL |
| K8 | 008-801-2177-G11 | QDATGVTPFD LAAYYGNEDI AEVLQKAAKL |
| K9 | 008-801-2178-B4 | QDKFGKTAFD ISIDNGNEDI AEVLQKAAKL |
| K10 | 008-801-2178-C11 | QDKWGTTPFD LAALIGNEDI AEVLQKAAKL |
| K11 | 008-801-2178-E11 | QDHMGETPFD LAAYFGNEDI AEVLQKAAKL |
| K12 | 008-801-2179-E4 | QDKFGKTPFD LAIDNGNEDI AEVLQKAAKL |
| K15 | 008-801-2179-E12 | QDLRGQTPFD LAAWQGNEDI AEVLQKAAKL |

**Figure S5. Sequences of DARPins identified in this study.** Sequences of the DARPins, with name and clone ID indicated, aligned using ClustalO. Amino-acid differences are shaded in grey except for the single difference between K2 and K5 shaded in cyan. Repeat, N-cap and C-cap regions are indicated above the sequences.

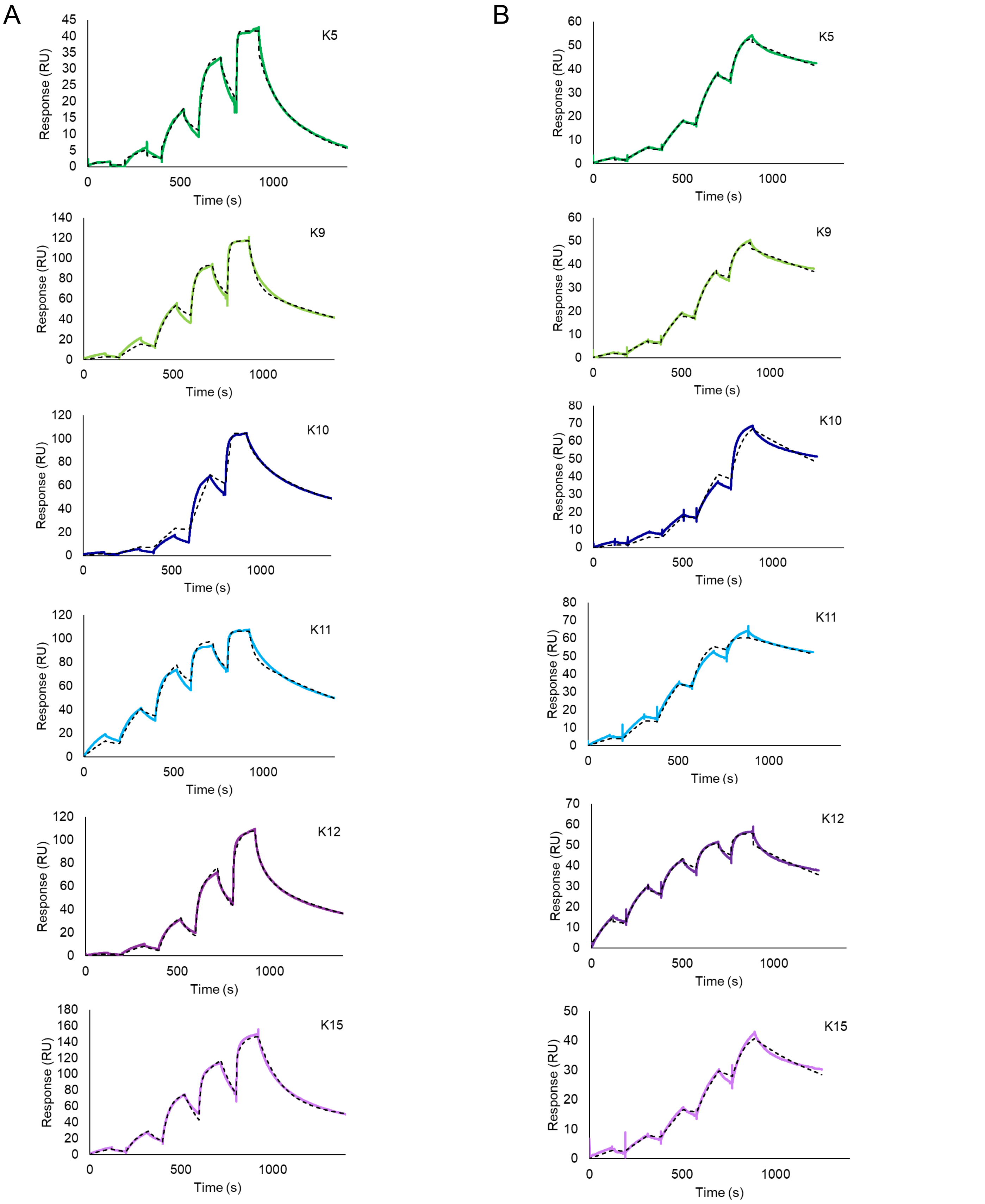


**Figure S6. Affinity of DARPin for CagI and CagI$^{C}$.** Surface Plasmon Resonance experiments using single-cycle-mode on CM5 chips coated with CagI (A) or with (B) CagI$^{C}$. DARPins were injected on the chips at increasing concentrations as follows. For experiments performed on full-length CagI, concentrations of DARPins were: 0.5, 2.5, 12.5, 62.5 and 312.5 nM for K9, K11, K12, K15; and 1, 3, 9, 27 and 81 nM for K10. For CagI$^{C}$ experiments, injections of DARPins were performed with concentrations of 0.05, 0.15, 0.45, 1.35 and 4 nM for K9, K10 and K11; 0.16, 0.8, 4, 20 and 100 nM for K12; and 2, 4, 8, 16 and 32 nM for K15. Fit curves obtained with heterogenous ligand model (CagI) or binding model 1:1 (CagI$^{C}$) are shown as black dashed lines.

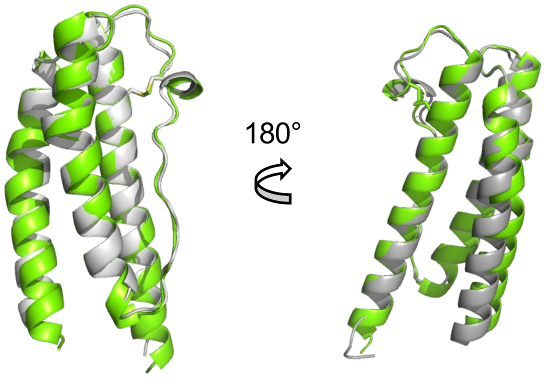


**Figure S7. Structural comparison of crystal structure of CagI$^{204\text{-}307}$ and corresponding region in the AF model.** Two rotated views of the crystal structure of CagI$^{204\text{-}307}$ from the CagI:K2 complex (green) and AF model (grey) displayed as cartoon. The side chains of the cysteine residues 272 and 283 forming the disulfide bond are displayed as ball-and-stick with sulfur atoms coloured in yellow.

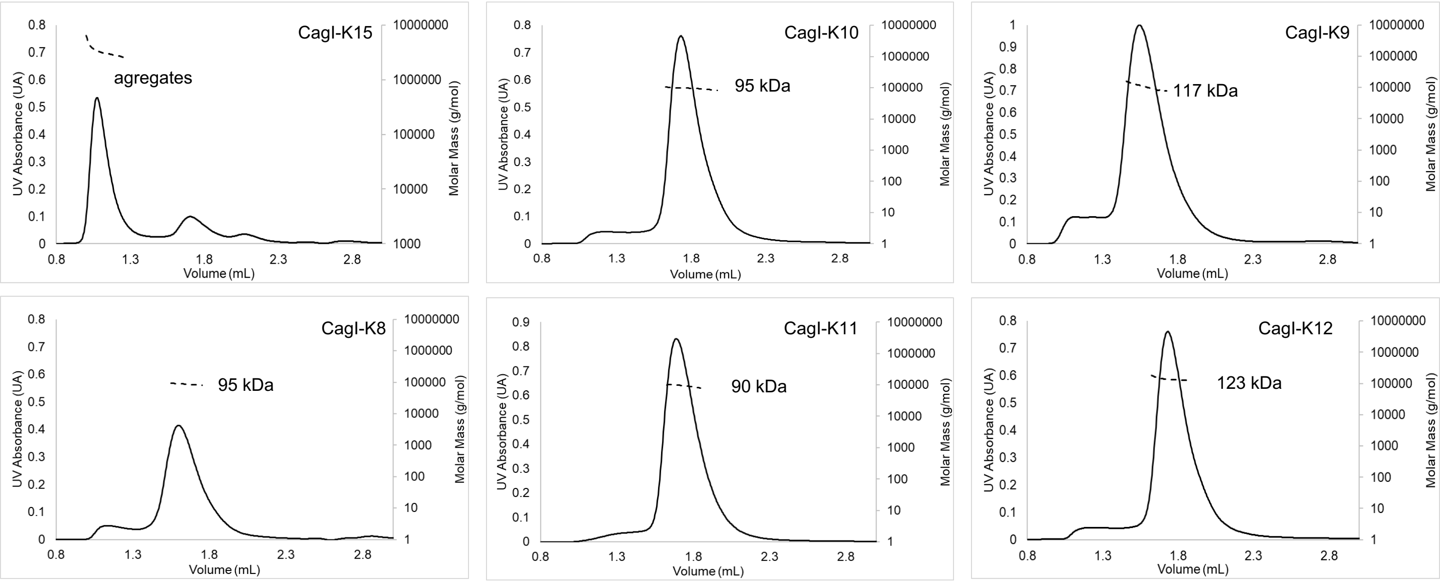


**Figure S8. Determination of molar mass of CagI:DARPins complexes by size exclusion chromatography coupled to multi-angle light scattering.** Each purified CagI:DARPin complex was submitted to SEC-MALS measurement represented by A280 chromatograms. Molar mass calculations are represented by dotted lines on each graph.

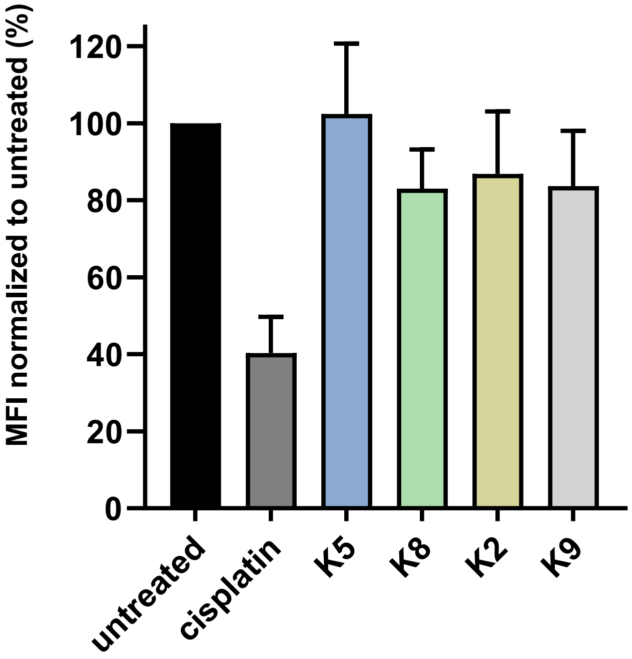


**Figure S9. Cell binding measurements by flow cytometry with *H. pylori* P12 [pHel12::gfp]** pre-treated with DARPins or left untreated. Data shown are mean values and standard deviations of median fluorescence intensities (MFI) normalized to untreated bacteria, resulting from three independent experiments.

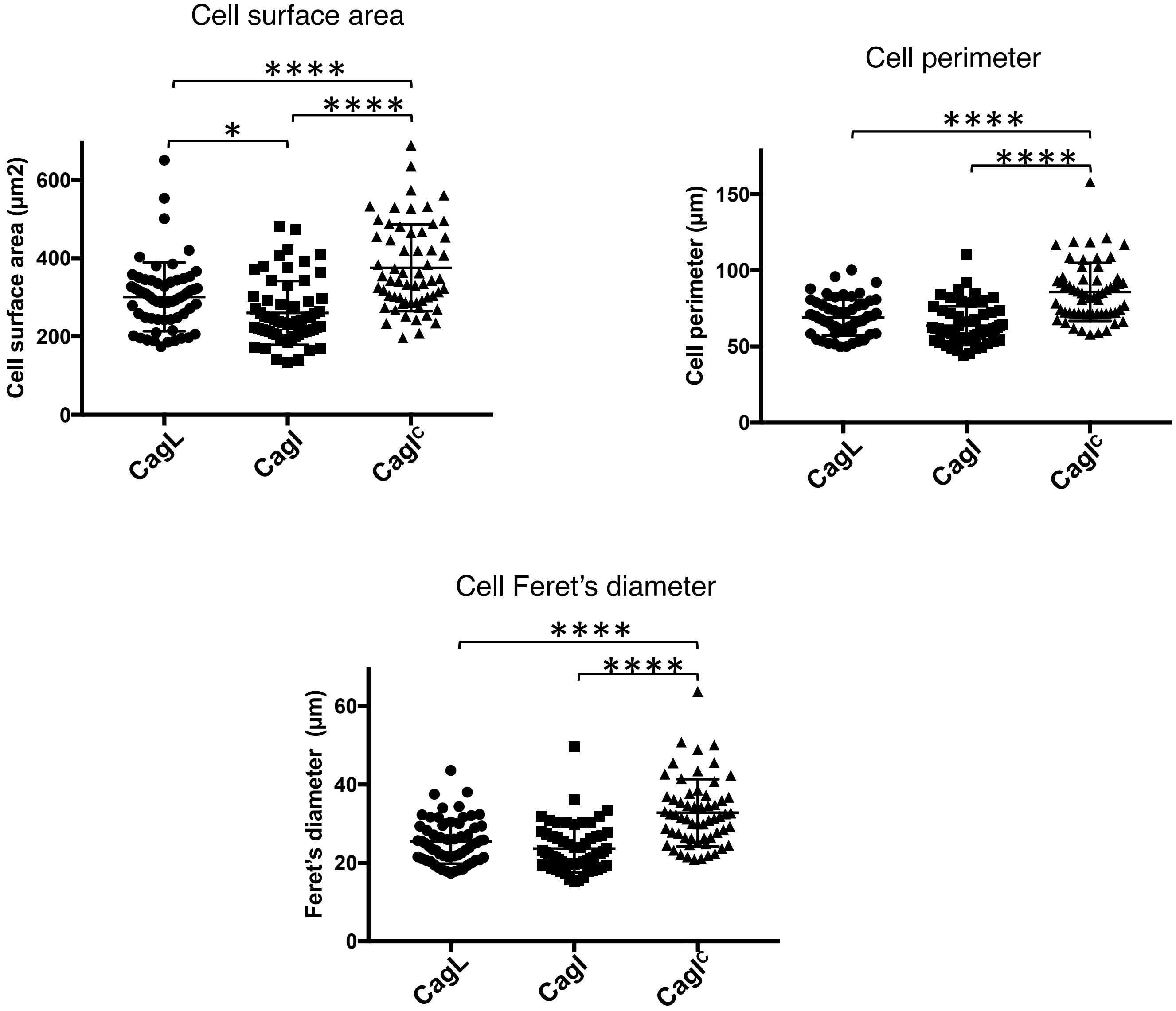


**Figure S10. Quantitative measurement of cell spreading on CagL, CagI or CagI$^C$.** Morphological characterization of cells adhered to 0.15 µg of CagL, CagI, CagI$^C$ 60 minutes after seeding. Cell surface area, perimeter and Feret's diameter were determined on phase contrast images using Fiji software. Each dot represents one cell (60 adhered AGS cells were measured for each condition). Each of the three parameters clearly shows that cells that have adhered to CagI$^C$ have a larger and more extended contact surface compared to cells bound to CagI or CagL. A one-way ANOVA with Tukey's post-test was used to determine significance. Means ± SD are shown with *p<0.1, ****p<0.0001.
